# Single Tube qPCR detection and quantitation of hotspot mutations down to 0.01% VAF

**DOI:** 10.1101/2021.08.12.456178

**Authors:** Kerou Zhang, Luis Rodriguez, Lauren Yuxuan Cheng, David Yu Zhang

## Abstract

Clinically and biologically, rare DNA sequence variants are significant and informative. However, existing detection technologies are either complex in workflow, or restricted in the limit of detection (LoD), or do not allow for multiplexing. Blocker displacement amplification (BDA) method can stably and effectively detect and enrich multiple rare variants with LoD around 0.1% variant allele fraction (VAF). Nonetheless, the detailed mutation information has to be identified by additional sequencing technologies. Here, we present allele-specific BDA (As-BDA), a method combining BDA with allele-specific TaqMan (As-TaqMan) probes for effective variant enrichment and simultaneous SNV profiling. We demonstrated that As-BDA could detect mutations down to 0.01% VAF. Further, As-BDA could detect up to four mutations with low to 0.1% VAF per reaction using only 15 ng DNA input. The median error of As-BDA in VAF determination is approximately 9.1%. Comparison experiments using As-BDA and droplet digital PCR (ddPCR) on peripheral blood mononuclear cell (PBMC) clinical samples showed 100% concordance for samples with mutations at ≥ 0.1% VAF. Hence, we have shown that As-BDA can achieve simultaneous enrichment and identification of multiple targeted mutations within the same reaction with high clinical sensitivity and specificity, thus helpful for clinical diagnosis.

## INTRODUCTION

Somatic mutations have been reported to play an essential role in many diseases, including cancer^1–5^. Somatic mutations within tumoral DNA are highly specific cancer biomarkers^3,4,6^, and the detection of which could be crucial and meaningful for diagnosis, prognosis, and precision medicine treatment. Traditionally invasive tissue biopsy, as the current gold standard for tumor diagnosis, has been reported to be less repeatable and less informative due to tumor heterogeneity. Also, tissue biopsy is sometimes hard to access in some types of tumors or in some patients with advanced disease^7–9^. Liquid biopsy is a non-invasive diagnostic technique that is capable of profiling tumor genomes with good concordance, including tests for circulating tumor DNA, circulating tumor cells or exosomes, and so on, in different fluids^10^. Moreover, liquid biopsy is helpful in establishing longitudinal tumor profiling of patients with more consistency. However, somatic mutations detected from the liquid biopsy usually have lower variant allele fraction (VAF), especially in early-stage patients. It has been challenging for current molecular diagnostics methods to achieve sensitive detection, efficient enrichment, and direct profiling of rare variants simultaneously while using readily accessible instruments. Allele-detection methods like amplification refractory mutation system (ARMS-PCR)^11^, allele-specific PCR (As-PCR)^12^, CAST-PCR^13,14^, microarray^15^ and deep-sequencing^16^, are limited either in plex number of detected mutations, or the limit of detection (LoD), or the complex and expensive experimental pipeline. Droplet digital PCR (ddPCR) can detect and visualize rare mutation with VAF low to 0.1%, while ddPCR is limited to single plex or duplex and a special set of ddPCR instruments is required. Furthermore, the dynamic range of DNA input for the ddPCR reaction is less flexible, as more input would lead to a higher chance of co-existence of more molecules within one droplet, influencing the accuracy of the result.

Blocker displacement amplification (BDA)^17^ method has been reported effective in enriching rare variant sequences with LoD as low as 0.1% VAF in multiplexed real-time PCR reactions. However, previous BDA applications in real-time PCR focused on applying TaqMan probes downstream to the blocker enrichment region to indicate the generation of the amplicon, thus non-specific to any mutations that happen within the blocker enrichment region. It requires at least two reactions^17^ or additional technology like Sanger sequencing^18^, next-generation sequencing (NGS)^19^ or third-generation sequencing^20^, to determine the identity of the enriched variant, increasing the total cost and overall turnaround time. Here we present an allele-specific BDA (As-BDA) method, which takes advantage of the effectiveness of BDA in rare variant enrichment and integrates allele-specific TaqMan probe for single nucleotide variant (SNV) genotyping. Using synthetic reference samples, we showed that our As-BDA method could detect rare variants low to 0.01% VAF, and simultaneously report the identity of the mutation within the same qPCR reaction by fluorescence signal from different channels. Then, we demonstrated that our As-BDA method could detect mutations with VAF down to 0.1% using only 15 ng DNA input, which is a low amount of DNA input that could be obtained in many sample types. Thus, As-BDA has reached the best LoD of current detection technologies could achieve with limited DNA amount, and As-BDA is also competitive in the detection of rare variants from 0.1% to 0.01% VAF, revealing the great advantage of As-BDA in detecting rare mutations over other current detection technologies including BDA. Further, we presented the multiplexed ability of As-BDA by designing an assay targeting the top 10 ranked mutations in *IDH2* within 3 tubes, with each tube could detect up to 4 mutations synchronously with as low as 0.1% VAF in 15 ng input. Moreover, we found successful VAF calling of commercially available reference samples from the threshold cycle (Ct) value by fitted regression equation from synthetic reference samples, with a median percent error around 9.1%. Lastly, we applied As-BDA in peripheral blood mononuclear cell (PBMC) samples from 27 acute myeloid leukemia (AML) patients and 24 healthy volunteers, and detected *IDH2* R140Q (419G>A) mutations in 3 AML patients. Additional comparative ddPCR analyses for R140Q mutations were performed on 11 AML samples and 7 healthy donor samples, and found 100% concordance among the results.

## MATERIALS AND METHODS

### Samples and study materials

Twenty-seven AML PBMC samples were purchased from Discovery Life Science (DLS). DLS collected these samples with written informed patient consent. Twenty-four PBMC samples from healthy volunteers were purchased from Zen-Bio, who collected the samples with written informed patient consent. IRB approval was not required as these are Exemption 4 (commercial, de-identified samples). Details of sample information are summarized in Tables S5 and S6.

### Oligonucleotides and repository samples

Primers, blockers, TaqMan probes and synthetic double-stranded DNA fragments (gBlocks) were purchased from Integrated DNA Technologies (IDT). Primers, blockers and synthetic double-stranded DNA fragments were purchased as standard desalting-purified, and TaqMan probes as HPLC-purified; they were resuspended in 1× IDTE buffer (10 mM Tris, 0.1 mM EDTA) in 100 uM (gBlocks were resuspended in 10 ng/μL) as the original stock and stored at 4 °C. Human cell-line gDNA NA18562 repository samples were purchased from Coriell Biorepository and stored at 4 °C for short-term usage, -20 °C for long-term storage. Commercial reference gDNA samples (Tru-Q 2 (5% Tier) Reference Standard (HD729), Tru-Q 7 (1.3% Tier) Reference Standard (HD734) and Myeloid DNA Reference Standard (HD829)) were purchased from Horizon Discovery and stored at 4 °C. Genomic DNA samples were further diluted with 1× IDTE buffer (IDT). GBlocks were serially diluted with 100ng/μL carrier RNA (Qiagen, 1017647) in 1× IDTE buffer with 0.1% TWEEN 20 (Sigma Aldrich).

### DNA extraction from PBMC samples

The DNA Blood Mini kit (Qiagen, 51104) was used on PBMC samples to extract DNA. NanoDrop spectrophotometer and Qubit Fluorometer were used to measure the yield of DNA. All DNA materials were stored at -20 °C until ready for analysis. As-BDA, ddPCR and NGS all used samples from the same DNA extract.

### As-BDA qPCR protocol

All As-BDA qPCR assays were performed on the CFX96 Touch Real-Time PCR Detection System using 96-well plates. PowerUp SYBR Green MasterMix (Thermo Fisher, A25742) was used for each qPCR reaction. For single-plex reactions, each 10 μL contains 200 nM *IDH2* R140 forward primer, 200 nM *IDH2* R140 reverse primer, 2 μM *IDH2* R140 BDA blocker and 200 nM As-TaqMan probe with 15 ng input DNA.

In tube 1, each reaction included 200 nM *IDH2* R140 forward primer, 200 nM *IDH2* R140 reverse primer, 2 μM *IDH2* R140 BDA blocker, 200 nM each As-TaqMan probes (four probes in total) and 15 ng input DNA per well with a final volume of 10 μL. For the reaction with 160ng DNA input, the final volume of each reaction is 20 μL with the concentrations of each component the same as described above.

In tube2 and 3, each reaction has 400 nM *IDH2* R172 forward primer, 400 nM *IDH2* R172 reverse primer, 4 μM *IDH2* R172 BDA blocker, 200 nM to 600 nM each As-TaqMan probes (details of composition concentration in Tables S1 to S3), as well as a set of 200 nM *GAPDH* forward primer, 200 nM *GAPDH* reverse primer, 200 nM *GAPDH* BDA blocker and 200 nM TaqMan probe targeting *GAPDH* housekeeping gene as internal control, and 15ng input DNA per well with a final volume of 10 μL. Reactions were performed in duplicate or triplicate. Details of design sequence are summarized in Tables S27 and S28.

The thermocycling program started with 3 min polymerase activation at 95 °C, followed by 60 repeated cycles of 10 s at 95 °C for DNA denaturing, 30 s at 60 °C for annealing/extension (95 °C:3 min – (95 °C:10 s – 60 °C:30s) × 60). Fluorescence signal was collected at 60 °C per cycle. All the reactions were conducted in duplicate or triplicate.

### Ct determination

Raw qPCR fluorescence data was processed in MATLAB. As described in previous research^17^, the average raw fluorescence signal from the first 10-15 cycles was treated as the background fluorescence signal and subtracted from each cycle. The post-subtraction fluorescence was regarded as final fluorescence. If the final fluorescence is lower than 200 relative fluorescence units (RFU), the well would be recognized as no amplification and of which the Ct value would be called as Inf. Else, the Ct value was termed by the interpolated fractional cycles at which reach the set threshold. Threshold was set different for each channel of each tube; details were shown in Table S4.

### Reference material preparation

Synthetic gBlocks were used to mimic real mutations. GBlocks were quantified by qPCR and diluted to approximately 1,500 molecules/μL for 100% VAF reference samples. The Ct values of the gBlocks were compared with and adjusted to the Ct values of 5 ng/μL genomic DNA (Coriell Institute, NA18562) assayed with the same primers. Then 100% VAF reference samples were mixed with 5 ng/μL wildtype (WT) genomic DNA to prepare reference samples of 10%, 5%, 1%, 0.5%, 0.3%, 0.1%, 0.05%, 0.03% and 0.01% by serial dilution. Reference samples with 10% VAF and 1% VAF were quantitated in NGS before being serially diluted to lower-VAF samples.

### *IDH2* R140Q ddPCR quantitation protocol

ddPCR mutation assay dHsaMDV2010057 (Bio-Rad, 10049550) was used to quantitate *IDH2* R140Q mutation and was performed on a QX200 Droplet Digital PCR System (Bio-Rad). A total of 20 μL reaction Master Mix containing 10 μL 2x ddPCR Supermix (Bio-Rad, 1863024), 1 μL of 20x target (FAM) and wildtype (HEX) primers/probe, and about 20ng of DNA sample was prepared and loaded onto the DG8 cartridges (Bio-Rad, 1864008). Then Droplet Generation Oil for Probes (Bio-Rad, 1963005) was added into another row of the DG8 cartridge, QX200 Droplet Generator (Bio-Rad, 10031907) was used to produce droplet emulsion. Then thermal cycling step started with 10 min at 95 °C, followed by 40 cycles of 30 s at 94 °C for DNA denaturation and1 min at 55 °C for annealing/extension. A 10 min incubation step at 98 °C was added to the end for enzyme deactivation, then the plate was held at 4 °C before the next step (abbreviated as 95 °C:10 min-(94 °C:30 s-60 °C:1 min) × 40-98 °C:10 min-4 °C:hold). After PCR, the plate was loaded onto the QX200 Droplet Reader (Bio-Rad, 1864003), and QuantaSoft software was used to collect droplet fluorescence data. Data analysis was conducted in MATLAB with code previously described^17^.

### NGS validation of VAF

We used NGS to validate our synthetic VAF templates and reference samples. We first used the same primer sets and PowerUp SYBR Green MasterMix (Thermo Fisher, A25742) as in the qPCR reactions to amplify the samples for 25 cycles; the thermocycling program started with 3 min polymerase activation at 95 °C, followed by 25 cycles of 10 s at 95 °C for DNA denaturing, 30 s at 60 °C for annealing/extension (95 °C:3 min – (95 °C:10 s – 60 °C:30s) × 25). Then we used end prep and ligation kit (New England Biolabs, E7546S for End Repair/dA-Tailing module, E7595S for Ligation module) or adaptor PCR [program: 95 °C:3 min – (95 °C:10 s – 60 °C:30s) × 2] to append adaptor sequence to the amplicons, followed by TruSeq index PCR amplification. We successfully validated both the 1% and 10% VAF of all the synthetic VAF templates.

## RESULTS

### Overview of allele-specific BDA (As-BDA) method

Here we presented the allele-specific BDA (As-BDA) method, where the designs of forward primers, reverse primers and blockers follow the same criteria as BDA^17^. We introduced the allele-specific TaqMan (As-TaqMan) probe into the BDA method, the sequence of which is designed to be partially overlapped with the 3’ region of the blocker to cover the target mutation, but not overlapped with forward primer (Figure 1A). Similar to BDA, the overlapped binding region between the blocker and the forward primer to the template sequence results in the competition in binding through a process of strand displacement. The standard free energy of forward primer displacing blocker is intentionally designed to be thermodynamically unfavorable in the case of a wildtype template. With a variant template, the blocker unfavorably binds to the variant template due to the energy penalty from the mismatch bubble or dangle formed between the blocker and variant template sequence. The energy penalty causes the kick-off of the blocker from the variant template by the forward primer, and the hybridization of the forward primer to the template. The binding energy of TaqMan probes is designed to be weaker than the blocker so that the TaqMan probes will not initially participate in strand displacement. Once a forward primer takes the place of a blocker and binds to a variant template, the corresponding As-TaqMan probe would bind to the variant and generate fluorescence readout during the amplification. Our As-BDA method is capable of detecting 0.1% VAF with 15ng input, where are only around 4 to 5 copies of the target variant within the reaction (Figure 1B), demonstrating the effective enrichment of variant and the sensitivity of As-TaqMan probes. As the ratio of wildtype to variant increases, there would be a much higher demand for the enrichment effectiveness of BDA to selectively amplify even the same number of variant molecules over the more wildtype molecules. Here, we showed that As-BDA could detect mutations with as low as 0.01% VAF using 160ng DNA (Figure 1C and Figure S2), which is roughly 10-fold more input, revealing the competitivity of As-BDA as we have not heard any other qPCR methods could achieve LoD with 0.01% VAF at the same level of stability, specificity, and sensitivity to the best of our knowledge. Sequences of amplification products were confirmed by Sanger Sequencing, as shown in Figure S17. Furthermore, our As-BDA method is 100% specific to the target mutation (Figure 1D). Upon the absence of the blocker, the As-TaqMan probe could only reach 10% LoD (Figure 1E), indicating the necessity of integration of As-TaqMan probes with BDA. The performance of As-BDA remained the same whether blocker and As-TaqMan probes were targeting the same strand (Figure 1C and 1D) or the opposite strands of a double-strand (Figure S1).

**Figure 1.**
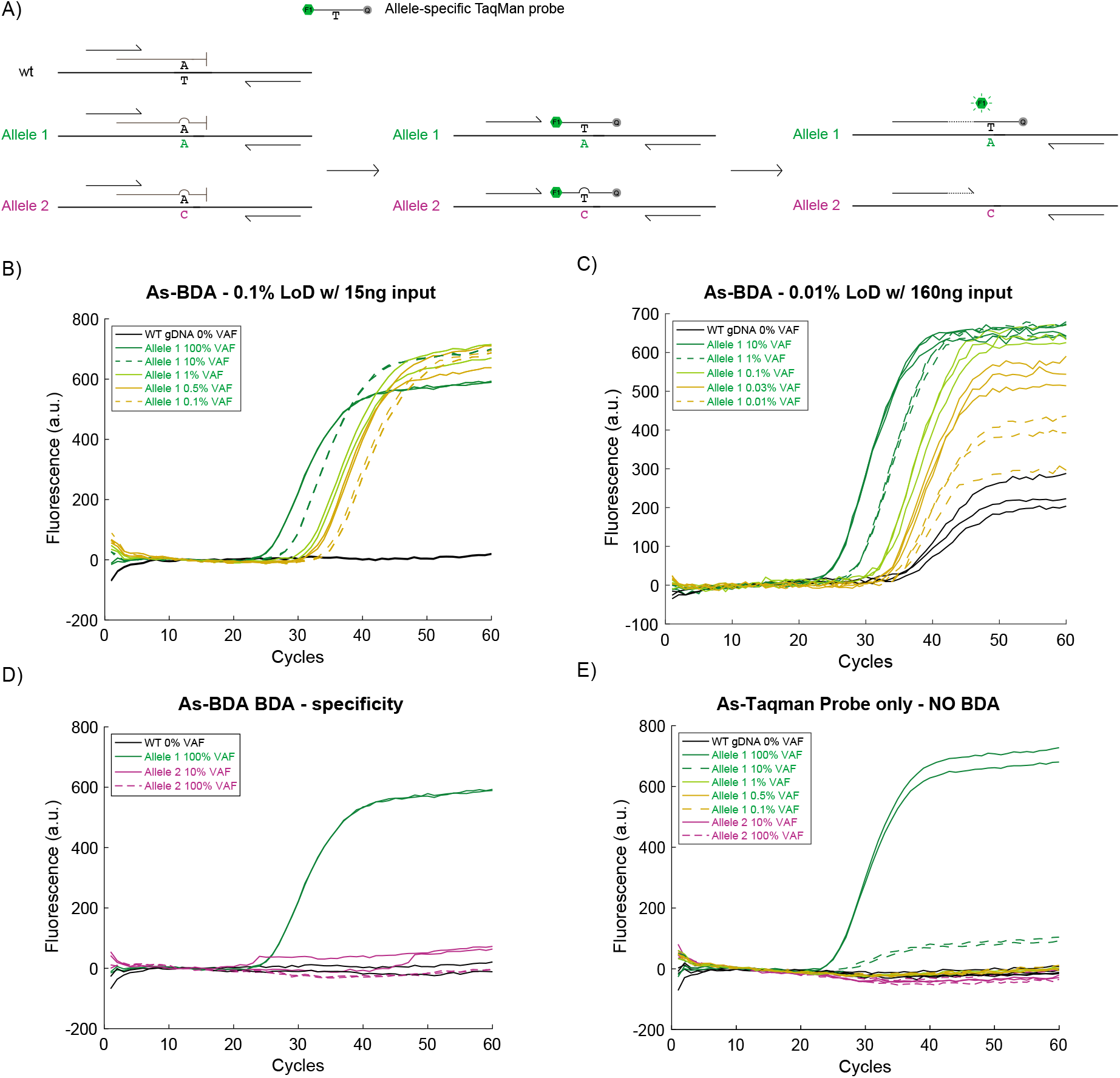
Overview of As-BDA. **(A). Schematic of As-BDA.** **(B). Sensitivity of As-BDA with 15ng input.** Single-plex result in *IDH2* 418 C>G mutation with As-BDA reveals the capability of detecting as low as 0.1% VAF with 15ng DNA input. Each reaction was performed in duplicate. **(C). Sensitivity of As-BDA with 160ng input.** The result of As-BDA in *IDH2* 418 C>T mutation reveals the capability of detecting as low as 0.01% VAF with 160ng DNA input, which is approximately 10-fold of 15ng, exhibits great advantage over LoDs of other current qPCR detection methods. Each reaction was performed in triplicate. **(D). Specificity of As-BDA.** The single-plex result with As-BDA shows clearly distinguishment of fluorescent signal between targeted variants versus non-targeted variants with 15ng DNA input, indicating the As-TaqMan probe is very specific to the target mutation. *IDH2* 418 C>G mutation is the target variant, *IDH2* 419 G>T is the ono-target variant. Each reaction was performed in duplicate. **(E). Performance of allele-specific TaqMan (As-TaqMan) probe.** Without BDA technology, the As-TaqMan probe could only effectively detect the existence of 10% VAF in *IDH2* 418 C>G mutation with 15ng DNA input, failing to detect lower-VAF samples. Each reaction was performed in duplicate.

### *IDH2* in AML and multiplexed As-BDA assay

*IDH2* is a gene encoding isocitrate dehydrogenase (NADP(+)) 2, an enzyme that catalyzes the oxidative decarboxylation of isocitrate to 2-oxoglutarate. Mutated *IDH2* would result in the production of oncometabolite (R)-2-hydroxyglutarate, promoting tumorigenesis like AML^21,22^, glioma^23^, and cholangiocarcinoma^24^. AML is an aggressive disease usually with a poor prognosis. There are about 20,240 new AML diagnosed cases, and an estimated 11,400 deaths every year in the United States, with the 5-year relative survival rate is only around 29.5%^25^. Per 100,000 men and women, there would be 4.3 cases of acute myeloid leukemia^25^. *IDH2* mutations usually happen more frequently within 8% ~ 19%^21,22,26^ AML patients compared with other cancers (Figure S3A). It has been reported that *IDH2* mutated patients had numerically worse 5-year overall survival and relapse-free survival^27^. Most *IDH2* mutations occur in the R140 and R172 loci (about 49% and 31%, respectively) (Figure S3B). Here, we applied As-BDA for multiplexed detection of the four most frequent R140 mutations and six most frequent R172 mutations in *IDH2* within three tubes (Table 1). Tube 1 targets four mutations in four separate fluorescent channels, excluding the FAM channel occupied by the internal integrated SYBR dye in PowerUp MasterMix. Each of tube 2 and tube 3 targets three mutations in three channels, with the Quasar channel functioning as the internal control channel in both tube 2 and tube 3 targeting a region of the housekeeping gene *GAPDH* to indicate the actual amount of sample input of each reaction.

**Table 1.** Details of *IDH2* assay design.

| Tube # | Position | Mutations | COSMIC ID | Nucleotide | Detection Channel |
| --- | --- | --- | --- | --- | --- |
| Tube1 | R140 | R140Q | 41590 | c.419G>A | Quasar |
|  |  | R140L | 41875 | c.419G>T | HEX |
|  |  | R140W | 41877 | c.418C>T | Cy5 |
|  |  | R140G | 1737874 | c.418C>G | Texas Red |
| Tube2 | R172 | R172S | 133672 | c.516G>C | Cy5 |
|  |  | R172M | 33732 | c.515G>T | Texas Red |
|  |  | R172W | 34039 | c.514A>T | HEX |
| Tube3 | R172 | R172S | 34090 | c.516G>T | Cy5 |
|  |  | R172K | 33733 | c.515G>A | Texas Red |
|  |  | R172G | 33731 | c.514A>G | HEX |

### Analytical assay validation

To evaluate the performance of our assay, we used synthetic gBlocks spiked in with wildtype (WT) genomic DNA to mimic samples with a gradient of VAFs. With 15 ng DNA input per tube, our assay detected 0.1% VAF for all mutations (Figure 2 and Figure S4 to S6). Moreover, the assay has no crosstalk channel fluorescence among any target mutation, indicating satisfying specificity.

**Figure 2.**
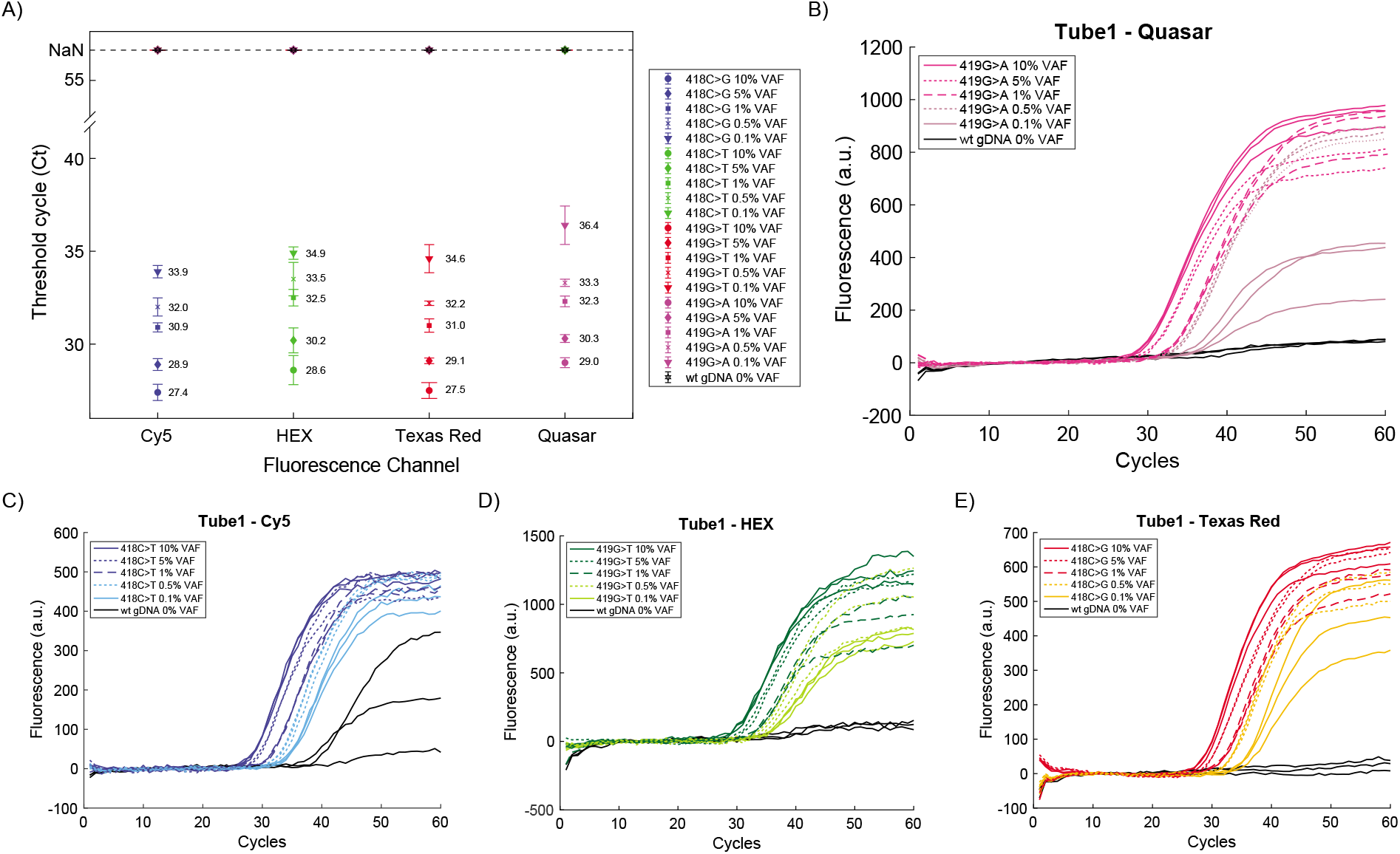
Performance of multiplexed As-BDA. **(A). Analytical summary of the Ct values in each channel of tube 1.** The Ct value of each mutation in each channel was collected and plotted, the error bar shows one standard deviation of each triplicate reaction. **(B-E). The qPCR curves of each channel.** For the triplicate reaction of WT gDNA in Cy5 channel as shown in **C)**, there are 2 out of 3 curves showing pop-up of fluorescence signal. Note that, only one curve has final fluorescence over 200 fluorescence units, so the final Ct called for the WT gDNA group is Inf. Each reaction was performed in triplicates, except the 5 % VAF groups were performed in duplicate.

For each variant of tube 1, reference samples with VAFs of 10%, 5%, 1%, 0.5% and 0.1% were assayed and the Ct value of each sample from all channels was collected based on the Ct determination criteria described in the method section. As shown in Figure 2A, Ct values from effective amplification were only manifest in fluorescent channels that are specific to the target mutations, with wildtype and all the non-target mutations revealed no Ct value and signals, indicating the high specificity of the assay in tube 1. The clear separation of the Ct values of each different VAF sample shown in Figure 2A demonstrates the capability of the assay to detect VAF down to 0.1% in every channel. qPCR curves of each channel of tube 1 are presented in Figure 2B to 2E. The absence of curves from most of wildtype groups indicates a low chance of false positives. We then validated the enrichment of each 0.1% VAF sample in Sanger sequencing (Figure S14), confirming the success of enriching 0.1% VAF samples and the trustworthiness of As-BDA qPCR results.

We continued to explore the performance of tube 2 and tube 3 by running samples from 100%, 10%, 5%, 1%, 0.3% to 0.1% VAF of each tube. The Ct distributions of tubes 2 and 3 are shown in Figures S4 and S5, respectively. Tubes 2 and 3 show similarly high specificity as that of tube 1, with neither cross-channel signal from any of the samples nor false-positive signals from wildtype samples. The qPCR curves shown in Figure S6 demonstrated the capability of our assay to detect each mutation with low to 0.1% VAF, the results of Sanger sequencing are shown in Figures S15, S16 and Table S26 accordingly.

### Assay validation with commercial reference samples

We further validated our assay using commercial reference samples. Based on the results from synthetic samples, we plotted Ct values of each duplicate or triplicate as individual dots versus logarithmic VAF values, from which a fitted relation was generated from linear regression. Ct values and log VAFs exhibited a strong linear correlation with an r-square of 0.991 for 419G>A mutation in tube 1 (Figure 3A) and the r-square is 0.986 for 515G>A mutation in tube 3 (Figure 3B). Fitting results of all the other mutations of the three tubes are shown in Figures S7 to S9. The fitted equation was used to quantitate the VAF of the unknown sample with the observed Ct value. We then used the 15 ng reference sample as input per reaction and compared the VAFs calculated from the obtained Ct value with those reported by the commercial vendor. Horizon reference samples Tru-Q 7 (1.3 % Tier) Reference Standard and Tru-Q 2 (5 % Tier) Reference Standard claimed 1.30 % and 5.0 % VAF of mutation 419G>A of tube 1. We ran As-BDA on these samples and calculated the VAFs, which are 1.69% and 5.50% as presented in Figure 3A. The closeness of the reference samples (shown as light blue star and magenta star in Figure 3A) to the fitting line exhibits the accuracy of the fitting result. The percentage error between the called VAF and claimed VAF is 30.0 % and 10.0 %. We then applied the same procedure on reference samples Tru-Q 7 (1.3 % Tier) Reference Standard and Myeloid DNA Reference Standard, which claimed 1.30 % and 5.00 % VAF for 515G>A in tube 3 (Figure 3B), with the percentage error between 3.8 % to 8.2 %. Results from Sanger sequencing are shown in Figure 3C and 3D to authenticate the results (Figure 3C and 3D).

**Figure 3.**
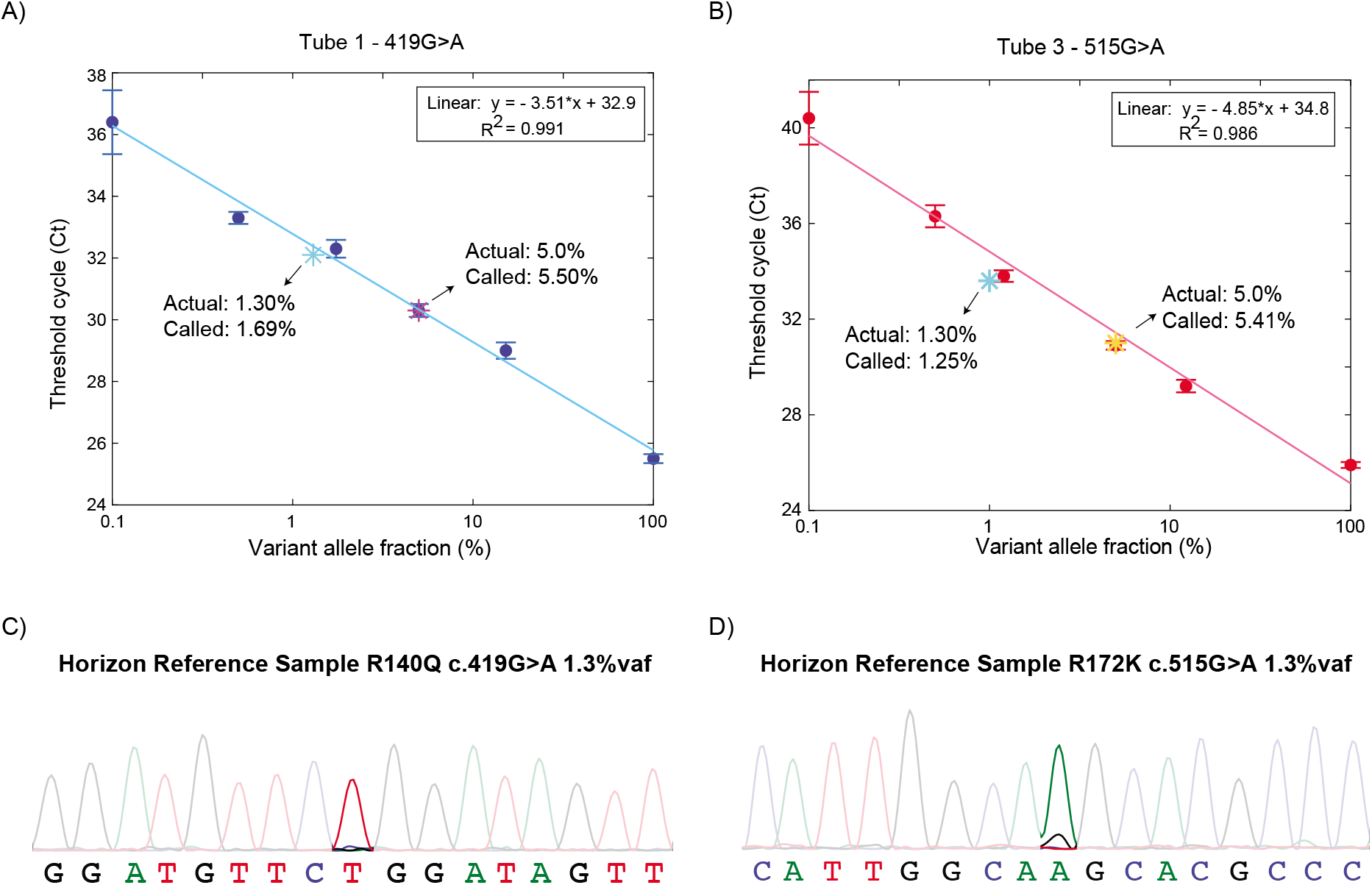
Validation of Ct determination with commercial reference samples. **(A). Standard VAF curve of the 419G>A mutation in tube 1.** The Ct values of the reference samples were plotted as individual blue dots; the fitted line is blue. The error bar shows one standard deviation. The 1.30% VAF and 5.0% reference samples were marked as a light blue star and a magenta star, respectively; their coordinates were determined by their actual VAF and the Ct value from the As-BDA reaction. The percent errors are 30% and 10%, respectively. **(B). Standard VAF curve of 515G>A mutation in tube 3.** The Ct values of the reference samples were plotted as individual red dots, the fitted line is magenta. The error bar shows one standard deviation. The 1.30% VAF and 5.0% reference samples were marked as a light blue star and a yellow star; their coordinates were determined by their actual VAF and the Ct value from the As-BDA reaction. The percent errors are 3.8% and 8.2%, respectively. **(C-D). Sanger sequencing validation of 1.3% reference sample.** Sanger sequencing curves show the enrichment of A allele versus G allele, the Sanger sequencing primer is the same reverse primer as that used in As-BDA qPCR reactions. Thus, the Sanger sequencing curves show the sequence of the reverse complementary strand.

### Clinical sample analysis

We performed As-BDA on the DNA extracted from PBMC samples of AML patients and healthy individuals. The Ct values of all samples from all the four channels are summarized in Tables S8 to S10 for tube 1, Tables S16 to S19 for tube 2 and tube 3. ddPCR using commercially available *IDH2* R140Q assay was performed 11 AML clinical samples, 7 healthy donor samples, 2 commercial reference samples, 1 synthetic reference material and the wildtype sample as shown in Figures S10 to S12 and Table S7. NGS was performed on 3 AML clinical samples, 2 commercial reference samples, 2 synthetic reference materials and wildtype sample targeting R140 region (Tables S12 to S15). For R172 region, 2 commercial reference samples, 2 synthetic reference materials and the wildtype sample were tested on NGS (Tables S20 to S25).

Using As-BDA, we found that 3 of the 27 clinical samples had fluorescent signals in the Quasar channel of tube 1, indicating the existence of *IDH2* R140Q mutations. VAFs of each clinical sample were then called based on the fitting equation. In summary, we performed ddPCR and As-BDA qPCR together for R140Q mutation on 11 AML clinical samples, 7 healthy donor samples as well as the reference samples. Here we compared results from As-BDA and ddPCR on each sample (Figure 4A). ddPCR called negative for the 3 AML clinical samples, 2 reference samples and 1 synthetic gBlock reference material, concordant as As-BDA results (Figure 4B-I). Comparison summary of the analytical result is shown in Table S11. These three double-confirmed AML clinical samples and the wildtype gDNA sample were later validated by Sanger sequencing (Figure S13). We repeated 1 AML clinical sample and 1 healthy donor sample each for nine times total in ddPCR and As-BDA qPCR reactions. All As-BDA qPCR results called negative for both the clinical sample and the healthy donor sample. For ddPCR, the clinical sample was totally called positive 1 out of 9 reactions, and the healthy donor sample was called positive 2 out of 9 reactions. The VAFs called positive by ddPCR are all around 0.1%. Thus, we hypothesize that the ddPCR could have false-positive results around its limit of detection, revealing the higher specificity of As-BDA compared with ddPCR.

**Figure 4.**
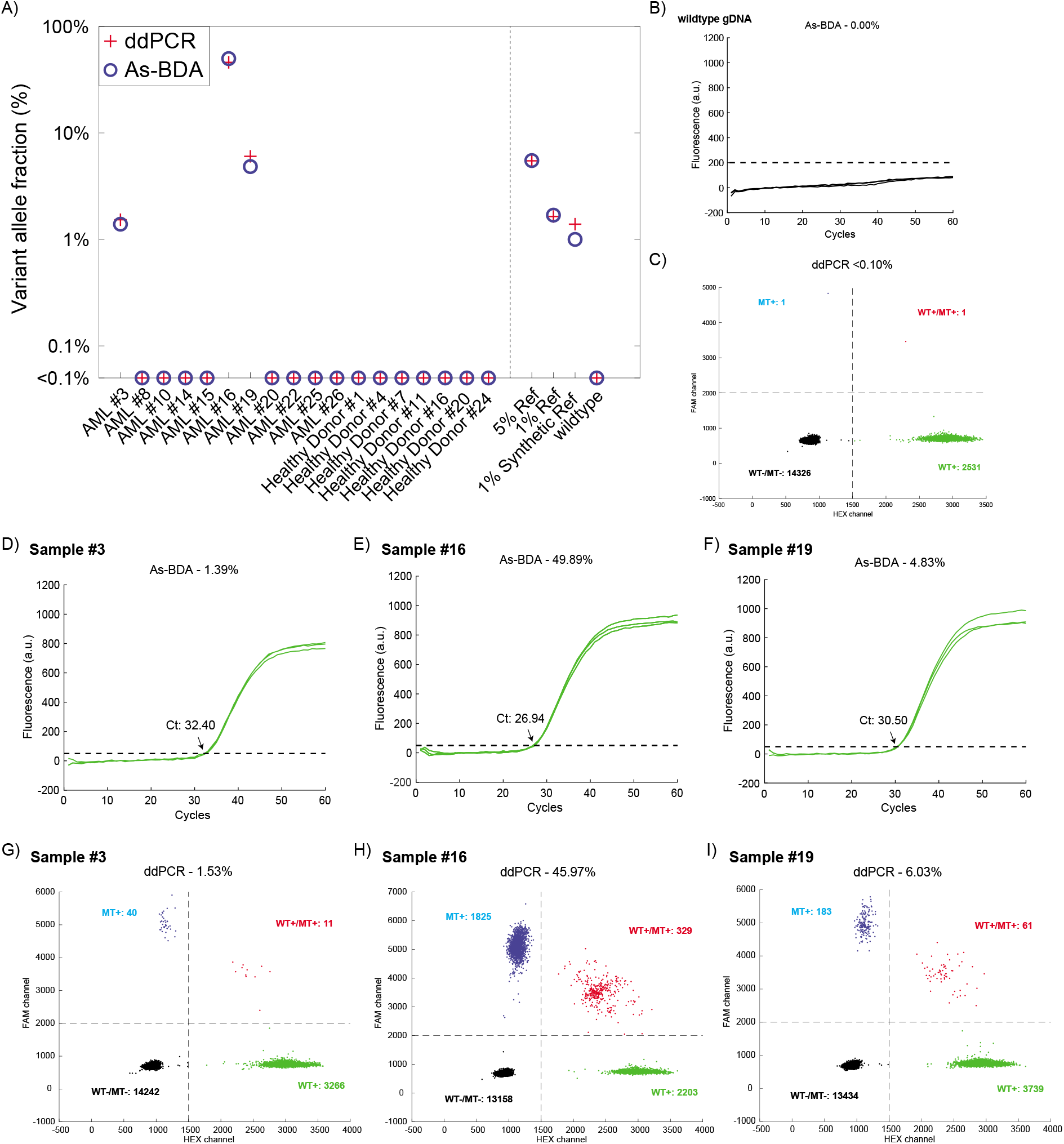
Comparative analytical results for clinical samples. **(A). Clinical sample summary of comparative analytical results in *IDH2* R140Q mutation.** Sample results are shown for the As-BDA reactions and the ddPCR assays. The DNA input was roughly 15 ng for each As-BDA reaction, and 20 ng for each ddPCR assay. **(B-C). As-BDA qPCR and ddPCR results of the wildtype sample.** **(D&G). As-BDA qPCR and ddPCR results of sample 3.** Green qPCR curves showed amplification of As-BDA as the threshold was plotted as the black dashed line. In ddPCR plots, green dots are HEX (wildtype) positive droplets, blue dots are FAM (variant) positive droplets, red dots are droplets containing both wildtype and variant templates, and black dots represent empty droplets. **(E&H). As-BDA qPCR and ddPCR results of sample 16.** **(F&I). As-BDA qPCR and ddPCR results of sample 19.**

## DISCUSSION

This As-BDA method presented here demonstrates integration with BDA and As-TaqMan probe in real-time PCR to allow simultaneous detection, enrichment and identification of multiplexed rare DNA mutations with a LoD low to 0.01% VAF inside the same reaction. This combination translates into practical advantages for molecular diagnostics and genomics research in at least two ways: detecting mutations qualitatively and quantitively at the same time, and enabling more rapid turnaround time of workflow, thus allowing result reporting within 2 hours, which are not currently achievable by ddPCR, NGS and other BDA methods.

The BDA technology takes advantage of blocker perfectly binding to the wildtype sequence, thus preventing wildtype sequences from being amplified and allowing variant sequences to be selectively amplified, resulting in the enrichment of variant sequences. However, the polymerase error introduced to the wildtype sequences during the initial thermocycling process, lead to the production of mutated wildtype amplicons and the generation of fluorescence signals. The fluorescent signal from mutation wildtype amplicons under the polymerase error rate hindered the indication of the existence of rare variant sequences at the similar magnitude level of VAF as the polymerase error, causing the limitation in LoD of BDA. With the integration of As-TaqMan probes, As-BDA could not only block the generation of wildtype amplicons using blocker, but also specifically indicate the production of target variant amplicons. In this way, our As-BDA is advantageous in the detection of VAF ≤ 0.1% VAF.

Real-time PCR instruments are broadly used in biolaboratory. Though ddPCR performs similarly in quantitation as As-BDA at the level of low DNA input, it requires special instrument sets, and has much lower flexibility in multiplexing. What is more, the dynamic input range of ddPCR is limited to less than 60 ng, as more input leads to a higher chance of co-existence of more molecules within a single droplet. Our As-BDA has much more flexibility in loading input, and we can even improve our sensitivity by the increase of input. Molecular NGS takes a longer time for the whole pipeline including library preparation, sequencing and data analysis, while diseases like AML are sensitive to the time of diagnosis, prognosis, and precision medicine treatment. To the best of our knowledge, we demonstrated we could detect up to 4 mutations inside a single tube qPCR reaction targeting mutations with VAF as low as 0.1% even 0.01% within two hours, which we did not see any other technology with this ability.

As-BDA method has been a proper candidate for non-invasive cancer diagnosis or monitoring based on liquid biopsy, As-BDA provides solutions for horizontally diagnosing cancer status and longitudinally monitoring dynamics of tumor genome in patients. In this study, we applied As-BDA to detect the 10 top-ranked mutations of *IDH2* R140 and R172 loci in PBMC samples from 27 AML patients and 24 healthy volunteers, and confirmed all the samples found R140Q mutation by Sanger sequencing. We believe our As-BDA method is a general method for detecting cancers and other diseases where rare variants play an important role. Also, our method could detect mutations with or even lower than 0.1% VAF using sample types like PBMC from the non-invasive collection, pointing out the potential of widespread application of As-BDA in clinical diagnosis and monitoring.

Although the current version of As-BDA has a relatively limited number of plexes, because the maximum number of useable fluorescence channels in qPCR instruments is only up to 6. We believe this is addressable by incorporating color combination strategy^29–31^ into the As-BDA method. In the future, we hope to upscale the plex capability of our assay to achieve more informative detection within the same reaction. Also, as As-BDA has shown great potential in detecting mutations down to 0.01% VAF, we would like to explore the further applications of As-BDA in detecting multiplexed mutations with VAF as low as 0.01% VAF and apply it to diseases like minimal residual disease (MRD).

## ACKNOWLEDGEMENTS

“The results shown in Figure S3A are in whole or part based upon data generated by the TCGA Research Network: https://www.cancer.gov/tcga.”

The authors thank Quoc-Khanh Pham and Dr. Bao for use of their ddPCR instrument.

The authors thank Dr. Paul Dolber, Jiaming Li, Xiangjiang Michael Wang and Xuwen Li for editorial assistance.

## AUTHOR CONTRIBUTIONS

K.Z and D.Y.Z conceived the project. K.Z and L.R performed the experiments and analyzed the data. K.Z and L.Y.C conducted As-BDA design. K.Z wrote the manuscript with input from all authors. D.Y.Z. revised the manusript.

## ADDITIONAL INFORMATION

Correspondence may be addressed to DYZ. DYZ declares a competing interest in the form of consulting for and significant equity ownership in NuProbe Global, Torus Biosystems and Pana Bio.

## DATA AVAILABILITY

The sequences of the DNA oligos and the oligo concentrations used for experiments are included in the supplementary file accompanying this manuscript. The main data supporting the results in this study are available within the paper and its Supplementary Information. The datasets collected and/or analyzed during the current study available from the corresponding author on reasonable request.

## CODE AVAILABILITY

The MATLAB code used for the ddPCR data analysis is available in the Supplementary Information.

## REFERENCES

1. Momozawa, Y. & Mizukami, K. Unique roles of rare variants in the genetics of complex diseases in humans. J. Hum. Genet. 66, 11–23 (2021).

2. Poduri, A., Evrony, G. D., Cai, X. & Walsh, C. A. Somatic mutation, genomic variation, and neurological disease. Science 341, 1237758 (2013).

3. Martincorena, I. & Campbell, P. J. Somatic mutation in cancer and normal cells. Science (80-.). 349, 1483 LP–1489 (2015).

4. Greenman, C. et al. Patterns of somatic mutation in human cancer genomes. Nature 446, 153–158 (2007).

5. Erickson, R. P. Somatic gene mutation and human disease other than cancer. Mutat. Res. 543, 125–136 (2003).

6. Watson, I. R., Takahashi, K., Futreal, P. A. & Chin, L. Emerging patterns of somatic mutations in cancer. Nat. Rev. Genet. 14, 703–718 (2013).

7. Cescon, D. W., Bratman, S. V, Chan, S. M. & Siu, L. L. Circulating tumor DNA and liquid biopsy in oncology. Nat. Cancer 1, 276–290 (2020).

8. Heitzer, E., Ulz, P. & Geigl, J. B. Circulating Tumor DNA as a Liquid Biopsy for Cancer. Clin. Chem. 61, 112–123 (2015).

9. Watanabe, K., Nakamura, Y. & Low, S.-K. Clinical implementation and current advancement of blood liquid biopsy in cancer. J. Hum. Genet. (2021) doi:10.1038/s10038-021-00939-5.

10. Remon, J. et al. Liquid biopsy in oncology: a consensus statement of the Spanish Society of Pathology and the Spanish Society of Medical Oncology. Clin. Transl. Oncol. 22, 823–834 (2020).

11. Newton, C. R. et al. Analysis of any point mutation in DNA. The amplification refractory mutation system (ARMS). Nucleic Acids Res. 17, 2503–2516 (1989).

12. Ugozzoli, L. & Wallace, R. B. Allele-specific polymerase chain reaction. Methods 2, 42–48 (1991).

13. Barbano, R. et al. Competitive allele-specific TaqMan PCR (Cast-PCR) is a sensitive, specific and fast method for BRAF V600 mutation detection in Melanoma patients. Sci. Rep. 5, 18592 (2015).

14. Didelot, A. et al. Competitive allele specific TaqMan PCR for KRAS, BRAF and EGFR mutation detection in clinical formalin fixed paraffin embedded samples. Exp. Mol. Pathol. 92, 275–280 (2012).

15. Ronald, J. et al. Simultaneous genotyping, gene-expression measurement, and detection of allele-specific expression with oligonucleotide arrays. Genome Res. 15, 284–291 (2005).

16. Kinde, I., Wu, J., Papadopoulos, N., Kinzler, K. W. & Vogelstein, B. Detection and quantification of rare mutations with massively parallel sequencing. Proc. Natl. Acad. Sci. 108, 9530 LP–9535 (2011).

17. Wu, L. R., Chen, S. X., Wu, Y., Patel, A. A. & Zhang, D. Y. Multiplexed enrichment of rare DNA variants via sequence-selective and temperature-robust amplification. Nat. Biomed. Eng. 1, 714–723 (2017).

18. Cheng, L. Y. et al. High sensitivity sanger sequencing detection of BRAF mutations in metastatic melanoma FFPE tissue specimens. Sci. Rep. 11, 9043 (2021).

19. Song, P. et al. Selective multiplexed enrichment for the detection and quantitation of low-fraction DNA variants via low-depth sequencing. Nat. Biomed. Eng. (2021) doi:10.1038/s41551-021-00713-0.

20. Thirunavukarasu, D. et al. Oncogene Concatenated Enriched Amplicon Nanopore Sequencing for Rapid, Accurate, and Affordable Somatic Mutation Detection. medRxiv 2020.11.12.20230169 (2020) doi:10.1101/2020.11.12.20230169.

21. Patel, K. P. et al. Acute myeloid leukemia with IDH1 or IDH2 mutation: frequency and clinicopathologic features. Am. J. Clin. Pathol. 135, 35–45 (2011).

22. Medeiros, B. C. et al. Isocitrate dehydrogenase mutations in myeloid malignancies. Leukemia 31, 272–281 (2017).

23. Yan, H. et al. IDH1 and IDH2 mutations in gliomas. N. Engl. J. Med. 360, 765–773 (2009).

24. Borger, D. R. et al. Frequent mutation of isocitrate dehydrogenase (IDH)1 and IDH2 in cholangiocarcinoma identified through broad-based tumor genotyping. Oncologist 17, 72–79 (2012).

25. Surveillance Research Program. SEER*Explorer: An interactive website for SEER cancer statistics. National Cancer Institute https://seer.cancer.gov/explorer/ (2021).

26. Papaemmanuil, E. et al. Genomic Classification and Prognosis in Acute Myeloid Leukemia. N. Engl. J. Med. 374, 2209–2221 (2016).

27. Ho, P. A. et al. Prevalence and Clinical Implications of IDH2 R140 and R172 Mutations in Older Adults with AML: A Report From SWOG,. Blood 118, 3516 (2011).

28. Stoler, N. & Nekrutenko, A. Sequencing error profiles of Illumina sequencing instruments. NAR Genomics Bioinforma. 3, (2021).

29. Xie, N. G. et al. High-throughput variant detection using a color-mixing strategy. bioRxiv 2021.06.30.450651 (2021) doi:10.1101/2021.06.30.450651.

30. Huang, Q. et al. Multicolor Combinatorial Probe Coding for Real-Time PCR. PLoS One 6, e16033 (2011).

31. Liao, Y. et al. Combination of fluorescence color and melting temperature as a two-dimensional label for homogeneous multiplex PCR detection. Nucleic Acids Res. 41, e76–e76 (2013).

