## Supplementary File for "Single Tube qPCR detection and quantitation of hotspot mutations down to 0.01% VAF"

**Section S1. Performance of As-BDA with reverse complementary As-TaqMan probe. 2**
**Section S2. Performance of As-BDA with 160ng DNA input. 3**
**Section S3. Overview of AML and *IDH2* gene. 4**
**Section S4. Details of 3-tube *IDH2* assay. 5**
**Section S5. Analytical assay performance. 7**
**Section S6. As-BDA, ddPCR and NGS results of PBMC samples from healthy donors and AML patients. 11**
**Section S7. Results of Sanger sequencing. 28**
**Section S8. List of As-BDA Oligonucleotide Sequences. 31**

### Section S1. Performance of As-BDA with reverse complementary As-TaqMan probe.

Compatibility of As-BDA with reverse complementary As-TaqMan probe was demonstrated and shown in Figure S1.

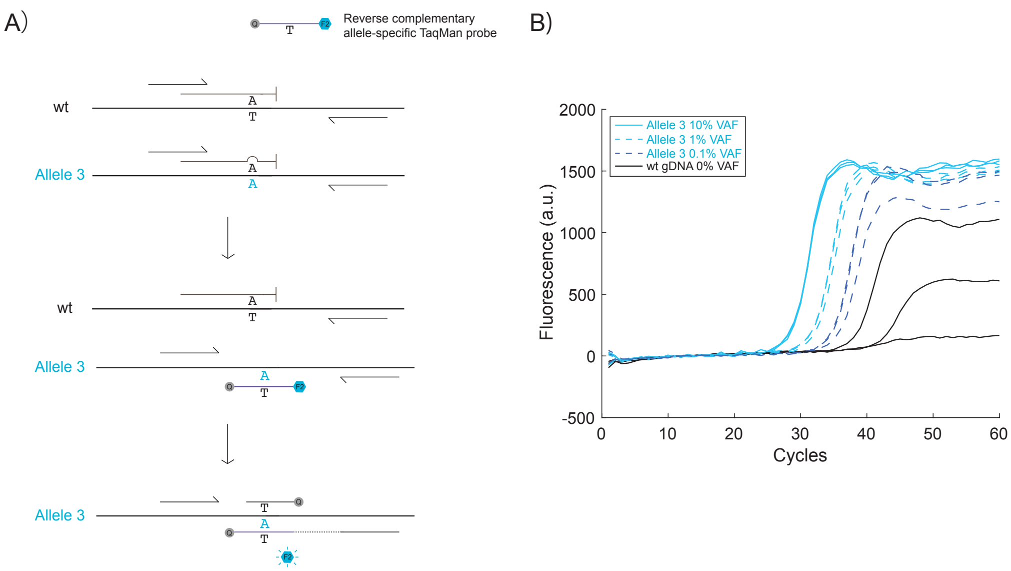


**Figure S1. Compatibility of As-BDA with reverse complementary As-TaqMan probe.**

**(A). Schematic of As-BDA with reverse complementary As-TaqMan probe.** The reverse complementary As-TaqMan probe could bind to the same strand as the reverse primer does. As the polymerase extends the reverse primer, if the reverse complementary As-TaqMan probe recognizes and binds to the template, it would be digested and then fluoresces.

**(B). Sensitivity of As-BDA with reverse complementary As-TaqMan probe.** The single-plex result of As-BDA in *IDH2* 418 C>T mutation is shown to have the LoD of 0.1% VAF with 15ng DNA input with reverse complementary allele-specific TaqMan probe. Each reaction was performed in triplicate.

**Section S2. Performance of As-BDA with 160ng DNA input.**

With 160ng DNA input, we demonstrated that As-BDA could detect VAF as low as 0.01% with a clear separation among each VAF samples and the wildtype sample in the qPCR curve, as shown in Figure S2.

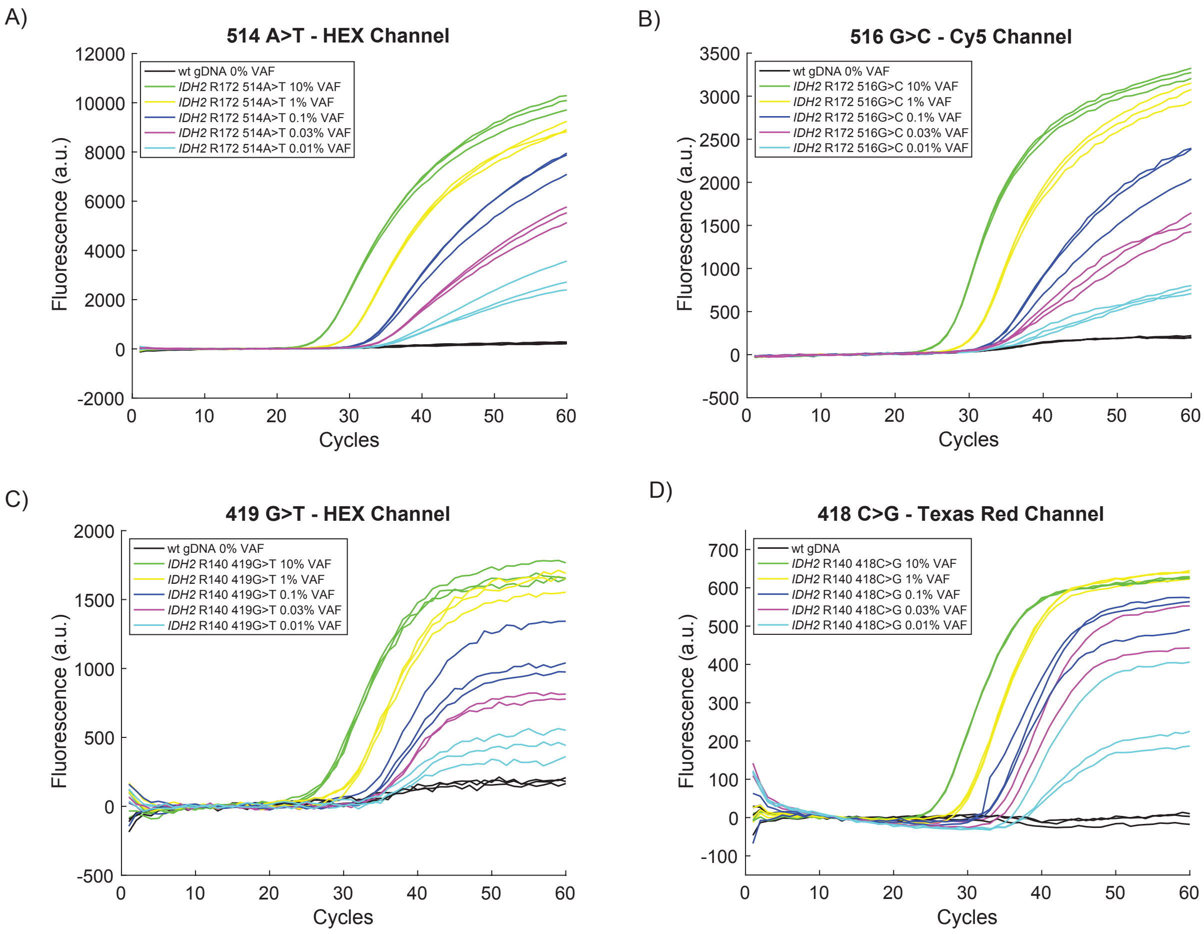


**Figure S2. Compatibility of As-BDA in detecting 0.01% VAF with 160ng DNA input per reaction.**

**(A). qPCR curves of As-BDA in detecting mutation 514 A>T from 10% VAF to 0.01% VAF in HEX channel.** Each reaction was performed in triplicate.

**(B). qPCR curves of As-BDA in detecting mutation 516 G>C from 10% VAF to 0.01% VAF in HEX channel.** Each reaction was performed in triplicate.

**(C). qPCR curves of As-BDA in detecting mutation 419 G>T from 10% VAF to 0.01% VAF in HEX channel.** Each reaction was performed in triplicate except the 0.03% VAF groups were performed in duplicate.

**(D). qPCR curves of As-BDA in detecting mutation 418 C>G from 10% VAF to 0.01% VAF in Texas Red channel.** Each reaction was performed in triplicate except the 0.03 % VAF groups were performed in duplicate.

### Section S3. Overview of AML and *IDH2* gene.

Based on the histogram distribution of *IDH2* alteration in different cancer, *IDH2* has the top 1 mutation frequency in Leukemia. Moreover, mutations happened within the *IDH2* R140 and R172 loci consist of 80% of the mutations in *IDH2*.

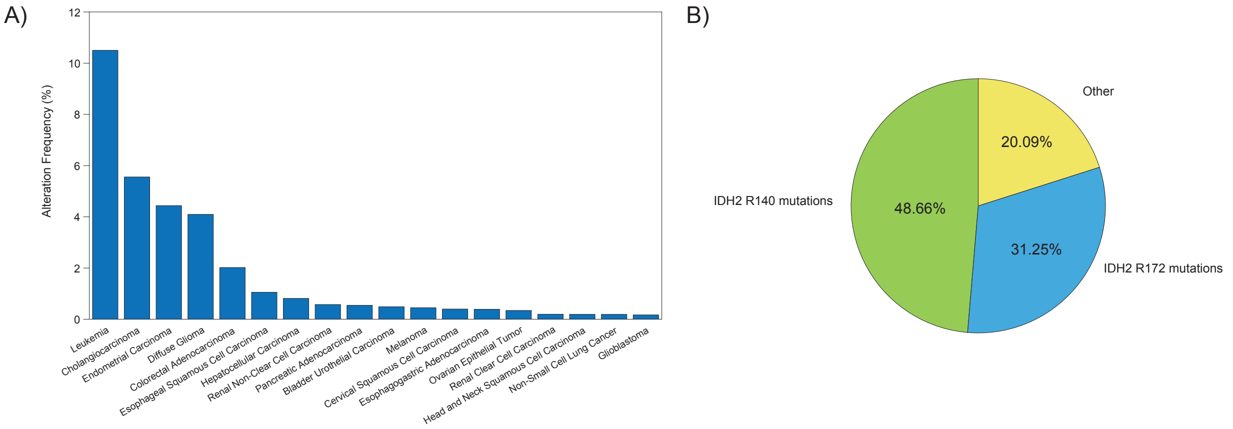


**Figure S3. Overview of AML and *IDH2* gene.**

**(A). Alteration frequency of *IDH2* in top-related cancers.** Based on TCGA PanCancer Atlas Studies, the histogram distribution of *IDH2* was plotted in the different cancers ranked by the mutation frequency.

**(B). *IDH2* mutation distribution.** Based on COSMIC[1] database, we collected all the reported *IDH2* mutations, and plotted the pie plot showing the high frequency of *IDH2* R140 and R172 in all *IDH2*-related mutations.

### Section S4. Details of 3-tube *IDH2* assay.

Detailed information regarding concentration of each component of each tube in As-BDA *IDH2* assay is provided in Table S1 to S3.

| Tube 1 | Stock concentration | Final concentration | Vol added (μL) |
|---|---|---|---|
| PowerUp MasterMix | 2X | 1X | 5 |
| *IDH2*_140_FP | 10 μM | 200 nM | 0.2 |
| *IDH2*_140_RP | 10 μM | 200 nM | 0.2 |
| *IDH2*_140_Blocker | 100 μM | 2000 nM | 0.2 |
| *IDH2*-140-419GT-TM | 5 μM | 200 nM | 0.4 |
| *IDH2*-140-418CT-TM | 5 μM | 200 nM | 0.4 |
| *IDH2*-140-418CG-TM | 5 μM | 200 nM | 0.4 |
| *IDH2*-140-419GA-TM | 5 μM | 200 nM | 0.4 |
| Template | 5 ng/μL | 15 ng | 3 |

**Table S1. Reaction mixture formulation for Tube 1.** The concentrations of each As-TaqMan probe are highlighted in light blue.

| Tube 2 | Stock concentration | Final concentration | Vol added (μL) |
|---|---|---|---|
| PowerUp MasterMix | 2X | 1X | 5 |
| *IDH2*_172_FP | 20 μM | 400 nM | 0.2 |
| *IDH2*_172_RP | 20 μM | 400 nM | 0.2 |
| *IDH2*_172_Blocker | 100 μM | 4000 nM | 0.4 |
| *IDH2*-172-516GC-TM | 10 μM | 200 nM | 0.2 |
| *IDH2*-172-515GT-TM | 10 μM | 200 nM | 0.2 |
| *IDH2*-172-514AT-TM | 10 μM | 400 nM | 0.4 |
| *GAPDH*_FP3 | 10 μM | 200 nM | 0.2 |
| *GAPDH*_RP3 | 10 μM | 200 nM | 0.2 |
| *GAPDH*_Blocker | 10 μM | 200 nM | 0.2 |
| *GAPDH*_TM | 10 μM | 200 nM | 0.2 |
| Template | 5 ng/μL | 15 ng | 3 |

**Table S2. Reaction mixture formulation for Tube 2.** The concentrations of each As-TaqMan probe are highlighted in light blue. The concentrations of the oligos targeting GAPDH housekeeping gene are highlighted in yellow.

| Tube 3 | Stock concentration | Final concentration | Vol added (μL) |
|---|---|---|---|
| PowerUp MasterMix | 2X | 1X | 5 |
| *IDH2*_172_FP | 20 μM | 400 nM | 0.2 |
| *IDH2*_172_RP | 20 μM | 400 nM | 0.2 |
| *IDH2*_172_Blocker | 100 μM | 4000 nM | 0.4 |

| | | | |
|---|---|---|---|
| *IDH2*-172-516GT-TM | 10 μM | 200 nM | 0.2 |
| *IDH2*-172-515GA-TM | 10 μM | 600 nM | 0.6 |
| *IDH2*-172-514AG-TM | 10 μM | 200 nM | 0.2 |
| *GAPDH*_FP3 | 10 μM | 200 nM | 0.2 |
| *GAPDH*_RP3 | 10 μM | 200 nM | 0.2 |
| *GAPDH*_Blocker | 10 μM | 200 nM | 0.2 |
| *GAPDH*_TM | 10 μM | 200 nM | 0.2 |
| Template | 5 ng/μL | 15 ng | 3 |

**Table S3. Reaction mixture formulation for Tube 3.** The concentrations of each As-TaqMan probe are highlighted in light blue. The concentrations of the oligos targeting GAPDH housekeeping gene are highlighted in yellow.

As described in the method section, thresholds were set different for each fluorescence channel of each tube for Ct value determination as shown in Table S4.

| Tube # | Cy5 (RFU) | HEX (RFU) | Texas Red (RFU) | Quasar (RFU) |
|---|---|---|---|---|
| 1 | 25 | 50 | 25 | 50 |
| 2 | 100 | 340 | 60 | 150 |
| 3 | 300 | 160 | 65 | 160 |

**Table S4. Threshold set for each channel of each tube.**

### Section S5. Analytical assay performance.

By running samples from 100%, 10%, 5%, 1%, 0.3% to 0.1% VAF of each tube, we collected and plotted the Ct distribution of tube 2 and tube 3 in Figure S4 and S5. Tube 2 and tube 3 show similarly high specificity as that of tube 1. There was neither cross-channel signal from any of the samples nor false-positive signal from the wildtype samples. The separation in Ct value of each VAF sample proved the ability of 0.1% VAF detection with 15 ng DNA input. Then qPCR curves of each channel of each tube were presents in Figure S6.

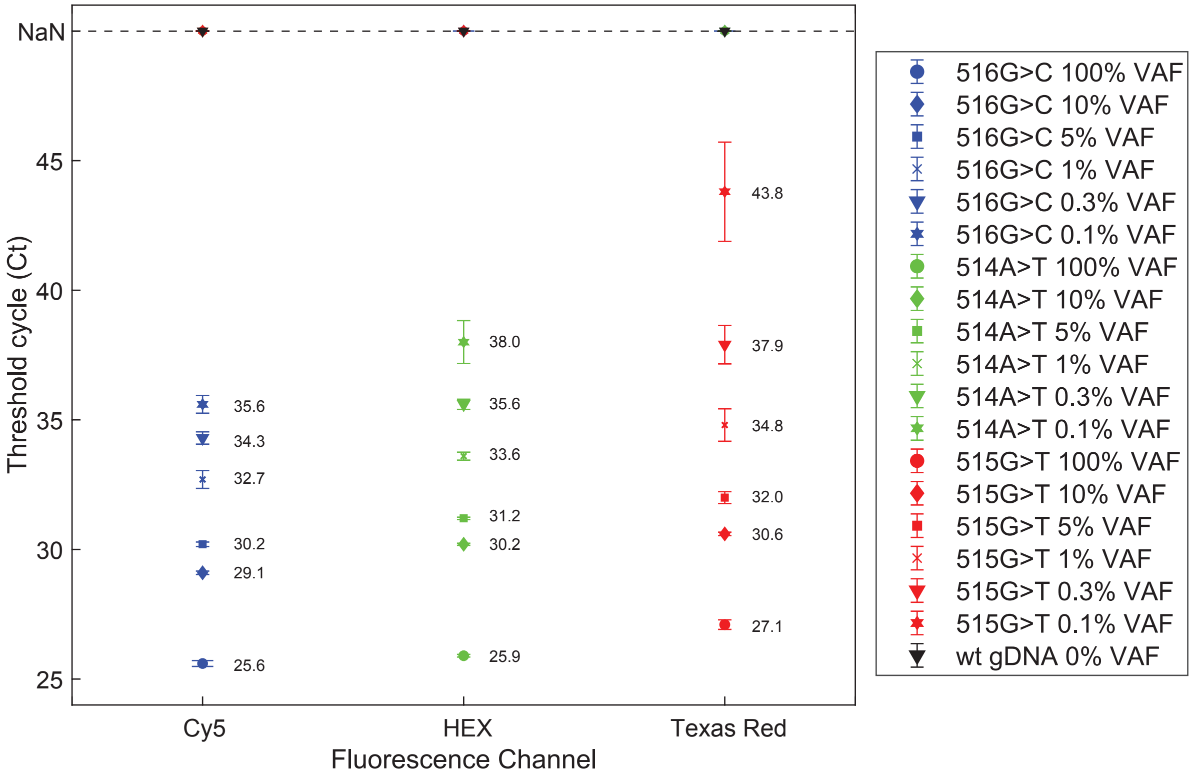


**Figure S4. Analytical summary of Ct value in each channel of tube 2.** Ct value of each mutation in each channel was collected and plotted, the error bar shows one standard deviation.

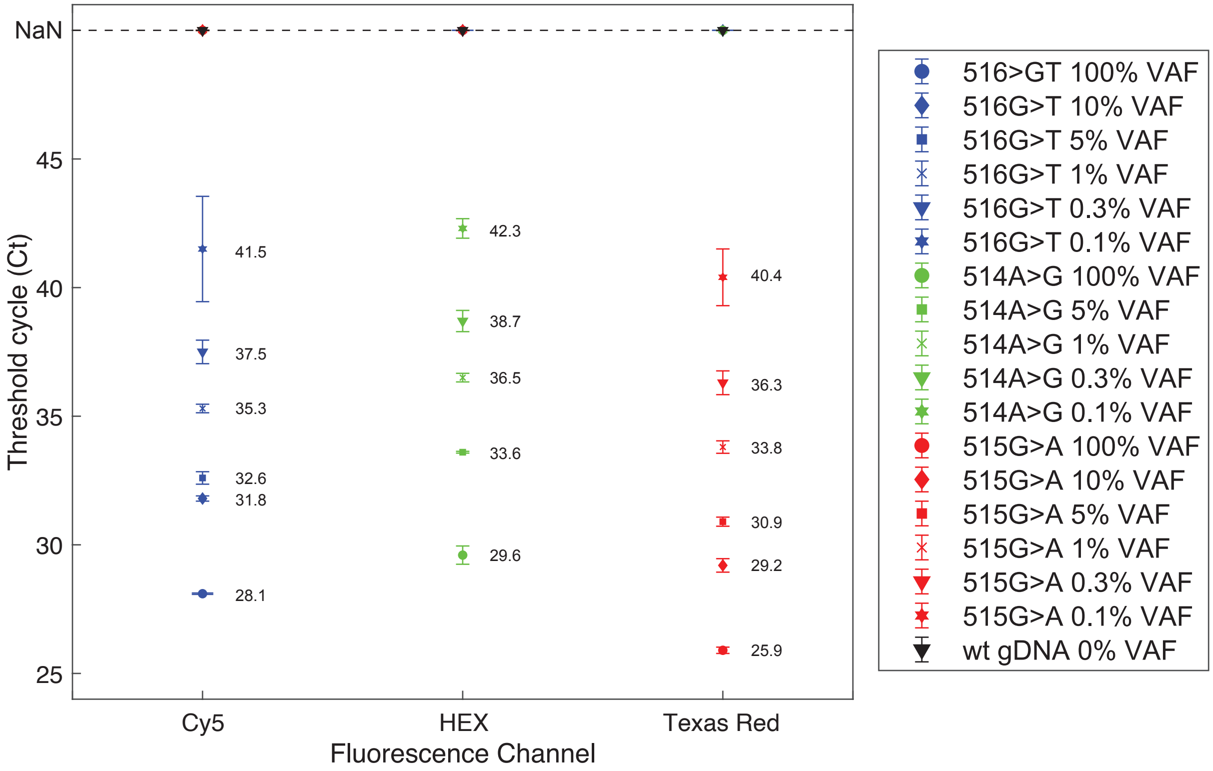


**Figure S5. Analytical summary of Ct value in each channel of tube 3.** Ct value of each mutation in each channel was collected and plotted, the error bar shows one standard deviation.

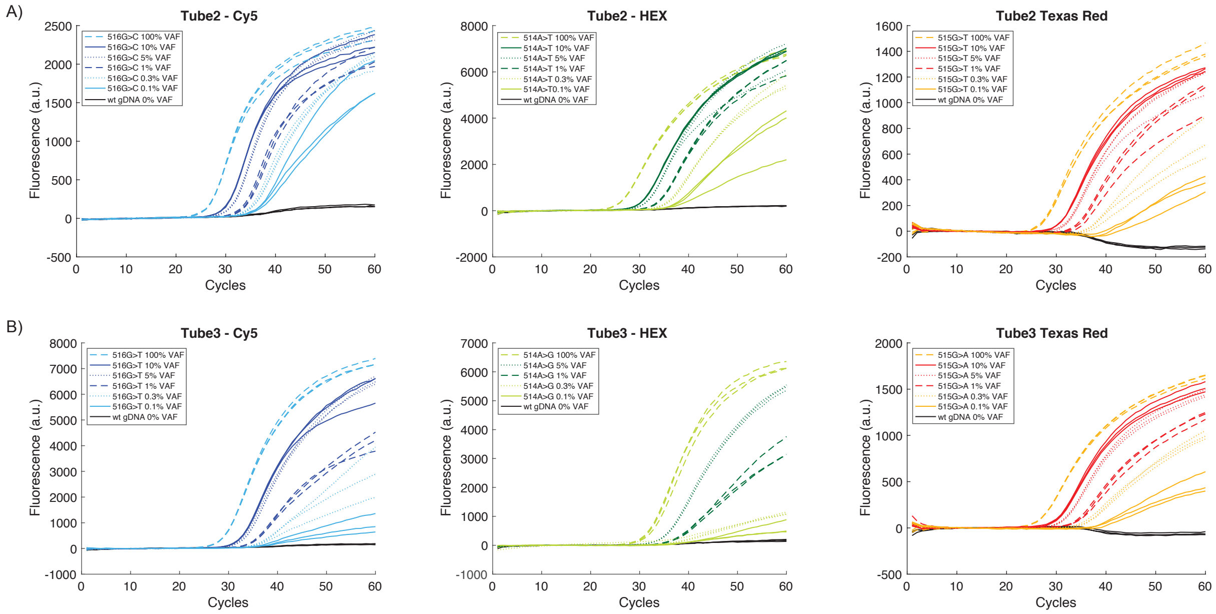


**Figure S6. qPCR curves of each channel of each tube.**

**(A). qPCR curves of tube 2.** Each reaction was performed in triplicates except the 5 % VAF groups were performed in duplicate.

**(B). qPCR curves of tube 3.** Each reaction was performed in triplicates except the 5 % VAF groups were performed in duplicate.

Ct values of each duplicate or triplicate reaction of each synthetic template were plotted versus logarithmic VAF values, as shown in Figure S7 to S9. Liner correlation was calculated based on curve fitting, providing formula for VAF quantitation.

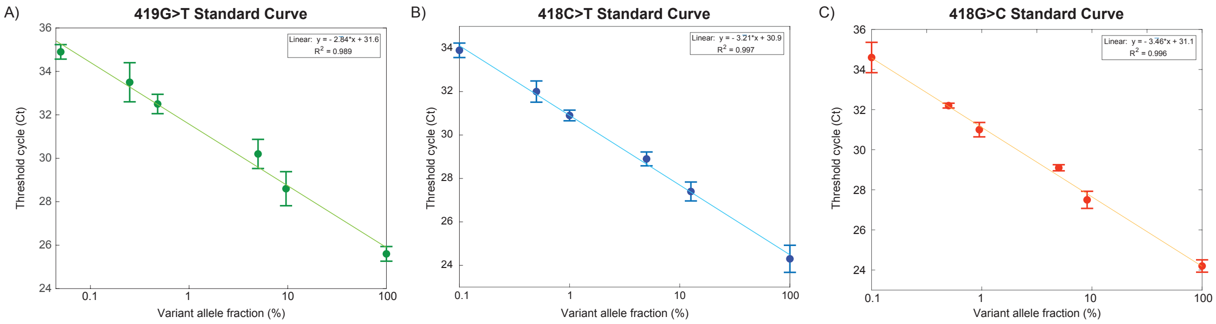


**Figure S7. Standard VAF curve of 3 mutations in tube 1.**

**(A). Standard VAF curve of 419G>T mutation in tube 1.** Triplicate or duplicate data points were plotted as individual dark green dots, the error bar shows one standard deviation. Fitted line was plotted as solid green. Ct values and log VAFs exhibited linear correlation with an $R^2$ is 0.989.

**(B). Standard VAF curve of 418C>T mutation in tube 1.** Triplicate or duplicate data points were plotted as individual dark blue dots, the error bar shows one standard deviation. Fitted line was plotted as solid blue. Ct values and log VAFs exhibited linear correlation with an $R^2$ is 0.997.

**(C). Standard VAF curve of 418G>C mutation in tube 1.** Triplicate or duplicate data points were plotted as individual red dots, the error bar shows one standard deviation. Fitted line was plotted as solid orange. Ct values and log VAFs exhibited linear correlation with an $R^2$ is 0.996.

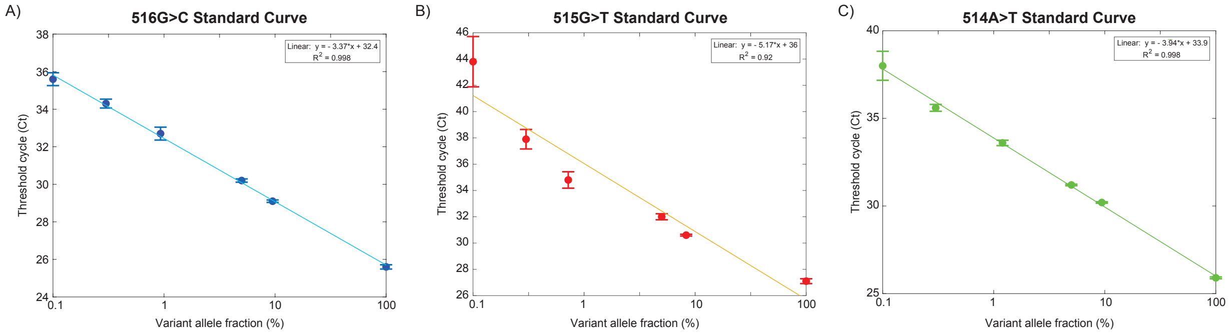


**Figure S8. Standard VAF curve of 3 mutations in tube 2.**

**(A). Standard VAF curve of 516G>C mutation in tube 2.** Triplicate or duplicate data points were plotted as individual dark blue dots, the error bar shows one standard deviation. Fitted line was plotted as solid blue. Ct values and log VAFs exhibited linear correlation with an $R^2$ is 0.998.

**(B). Standard VAF curve of 515G>T mutation in tube 2.** Triplicate or duplicate data points were plotted as individual red dots, the error bar shows one standard deviation. Fitted line was plotted as solid orange. Ct values and log VAFs exhibited linear correlation with an $R^2$ is 0.92.

**(C). Standard VAF curve of 514A>T mutation in tube 2.** Triplicate or duplicate data points were plotted as individual green dots, the error bar shows one standard deviation. Fitted line was plotted as solid green. Ct values and log VAFs exhibited linear correlation with an $R^2$ is 0.998.

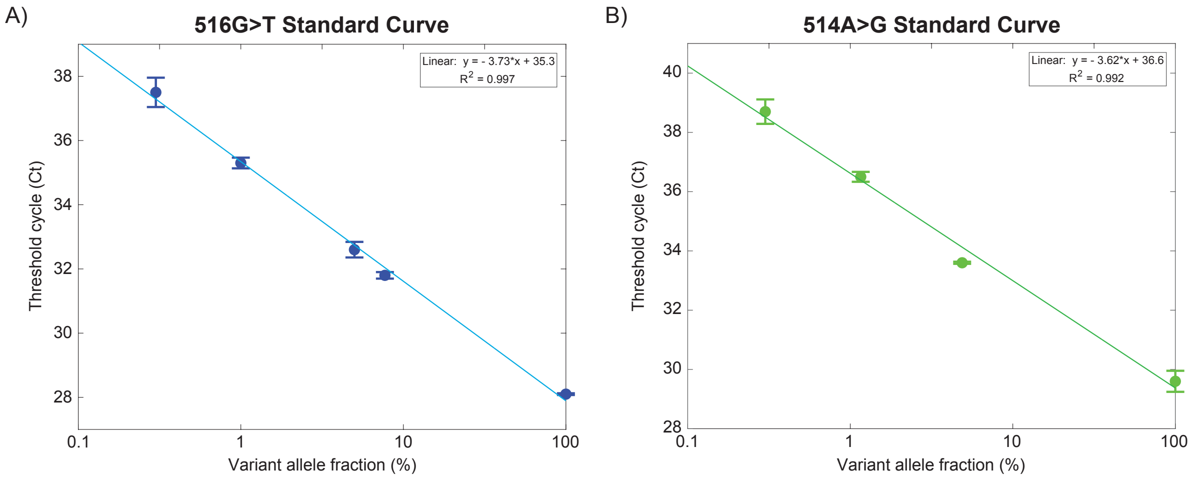


**Figure S9. Standard VAF curve of 2 mutations in tube 3.**

**(A). Standard VAF curve of 516G>T mutation in tube 3.** Triplicate or duplicate data points were plotted as individual dark blue dots, the error bar shows one standard deviation. Fitted line was plotted as solid blue. Ct values and log VAFs exhibited linear correlation with an $R^2$ is 0.997.

**(B). Standard VAF curve of 514A>G mutation in tube 3.** Triplicate or duplicate data points were plotted as individual green dots, the error bar shows one standard deviation. Fitted line was plotted as solid green. Ct values and log VAFs exhibited linear correlation with an $R^2$ is 0.992.

### Section S6. As-BDA, ddPCR and NGS results of PBMC samples from healthy donors and AML patients.

27 clinical AML PBMC samples were purchased from discovery life science and 24 healthy donor PBMC samples were from Zen-Bio, sample details are shown in Table S5 and S6.

| Sample ID | Patient ID | Primary Diagnosis | Disease status | Gender | Patient Age at Collection | Draw Date | Treatment Status | Clinical Stage |
|---|---|---|---|---|---|---|---|---|
| 1 | 121713346 | Leukemia, Acute Myeloid (AML) | Stable | Male | 82 | 7/20/20 | Active | / |
| 2 | 200000471 | Leukemia, Acute Myeloid (AML) | Stable | Male | 41 | 12/10/19 | Post | / |
| 3 | 121592979 | Leukemia, Acute Myeloid (AML) | Stable | Male | 91 | 5/4/20 | Active | / |
| 4 | 121582633 | Leukemia, Acute Myeloid (AML) | Stable | Female | 61 | 5/4/20 | Active | / |
| 5 | 200003029 | MDS/AML | Stable | Male | 67 | 3/20/19 | Post | / |
| 6 | 200008127 | Leukemia, Acute Myeloid (AML) | Newly Diagnosed | Female | 64 | 10/4/19 | Pre | M4 (FAB) |
| 7 | 200000471 | Leukemia, Acute Myeloid (AML) | Stable | Male | 41 | 12/10/19 | Post | / |
| 8 | 110025045 | Leukemia, Acute Myeloid (AML) | Stable | Female | 61 | 7/7/17 | Post | M1 |
| 9 | 200009066 | Leukemia, Acute Myeloid (AML) | Newly Diagnosed | Male | Unknown | 7/14/20 | Active | / |
| 10 | 200000340 | Leukemia, Acute Myeloid (AML) | Newly Diagnosed | Female | 38 | 5/15/19 | Pre | M4 |
| 11 | 200018300 | Leukemia, Acute Myeloid (AML) | Refractory | Male | 65 | 10/19/20 | Post | / |
| 12 | 200009196 | Leukemia, Acute Myeloid (AML) | Stable | Female | 58 | 2/19/20 | Active | / |
| 13 | 200013114 | Leukemia, Acute Myeloid (AML) | Unkown | Male | 83 | 10/29/19 | Pre | M5 |
| 14 | 200009609 | Leukemia, Acute Myeloid (AML) | Progressive | Female | 60 | 9/2/20 | Post | / |
| 15 | 200018824 | Leukemia, Acute Myeloid (AML) | Newly Diagnosed | Female | 56 | 1/14/21 | Pre | M4 |
| 16 | 200018595 | Leukemia, Acute Myeloid (AML) | Relapse | Male | 56 | 12/7/20 | Post | M4 |
| 17 | 200009007 | Leukemia, Acute Myeloid (AML) | Refractory | Male | 53 | 2/25/20 | Post | M4 |
| 18 | 200003963 | Leukemia, Acute Myeloid (AML) | Refractory | Female | 55 | 12/17/19 | Active | / |
| 19 | 200009536 | Leukemia, Acute Myeloid (AML) | Newly Diagnosed | Female | 61 | 5/4/20 | Pre | / |
| 20 | 200009469 | Leukemia, Acute Myeloid (AML) | Progressive | Male | 71 | 12/8/20 | Post | / |
| 21 | 200018487 | Leukemia, Acute Myeloid (AML) | Newly Diagnosed | Female | 49 | 12/22/20 | Pre | M5 |
| 22 | 200009002 | Leukemia, Acute Myeloid (AML) | Newly Diagnosed | Female | 76 | 2/11/20 | Post | AML (Blast crisis after CML) |

| | | | | | | | | |
|---|---|---|---|---|---|---|---|---|
| 23 | 121765714 | Leukemia, Acute Myeloid (AML) | Stable | Male | 73 | 11/5/20 | Active | / |
| 24 | 200013554 | Leukemia, Acute Myeloid (AML) | Stable | Female | 70 | 12/11/19 | Post | / |
| 25 | 200009189 | Leukemia, Acute Myeloid (AML) | Refractory | Female | 62 | 2/24/20 | Post | / |
| 26 | 200005227 | Leukemia, Acute Myeloid (AML) | Unkown | Male | 62 | 9/10/19 | Post | M0 |
| 27 | 121624482 | Leukemia, Acute Myeloid (AML) | Stable | Female | 32 | 7/20/20 | Active | / |

**Table S5. AML patient information including age at collection, gender, draw date, treatment status, disease status and clinical stage (optional).**

| Sample ID | Patient ID | Health Status | Gender | Patient Age at Collection | BMI | Diabetic | Smoker |
|---|---|---|---|---|---|---|---|
| 1 | PBMC093020C | Healthy | F | 64 | Unknown | Unknown | Unknown |
| 2 | PB000194 | Healthy | M | 21 | 31.5 | N | N |
| 3 | PBMC070218B | Healthy | M | 46 | 21.7 | N | N |
| 4 | PBMC070218A | Healthy | M | 47 | 25.1 | N | N |
| 5 | PBMC070218C | Healthy | F | 46 | 28.3 | N | N |
| 6 | PBMC012021B | Healthy | M | 56 | Unknown | Unknown | Unknown |
| 7 | PBMC012021C | Healthy | F | 73 | Unknown | Unknown | Unknown |
| 8 | PBMC012021D | Healthy | M | 73 | Unknown | Unknown | Unknown |
| 9 | PBMC120220F | Healthy | F | 69 | Unknown | Unknown | Unknown |
| 10 | PBMC120220A | Healthy | F | 65 | Unknown | Unknown | Unknown |
| 11 | PBMC111319B | Healthy | F | 38 | U | Unknown | Unknown |
| 12 | PBMC091520C | Healthy | M | 52 | 30.1 | N | N |
| 13 | PBMC091520B | Healthy | M | 35 | 25.1 | N | N |
| 14 | 00PB000415 | Healthy | M | 20 | 31 | N | N |
| 15 | PBMC061919B | Healthy | M | 54 | 28.7 | N | N |
| 16 | PBMC120920I | Healthy | F | 54 | Unknown | Unknown | Unknown |
| 17 | PBMC011520F | Healthy | M | 51 | Unknown | Unknown | Unknown |
| 18 | PBMC090220B | Healthy | M | 45 | Unknown | Unknown | Unknown |
| 19 | PBMC090220H | Healthy | F | 27 | Unknown | Unknown | Unknown |
| 20 | PBMC090120B | Healthy | M | 39 | 25 | Unknown | Y |
| 21 | PBMC090220F | Healthy | M | 58 | Unknown | Unknown | Unknown |
| 22 | PBMC090220D | Healthy | F | 56 | Unknown | Unknown | Unknown |
| 23 | PBMC090220C | Healthy | M | 59 | Unknown | Unknown | Unknown |
| 24 | PBMC091620B | Healthy | F | 64 | Unknown | Unknown | Unknown |

**Table S6. Healthy donor information including age at collection, gender, health status, BMI, diabetic status, smoking status.**

We applied As-BDA in 27 AML PBMC samples and 24 healthy PBMC samples, each sample was conducted in duplicate for each designed tube. The obtained Ct values and accordingly called VAFs of tube 1 were presented in Table S8 to S10. As-BDA reactions reported 3 AML patients with the R140Q (419 G>A) mutation, with other AML samples and all the healthy donor samples reported negative.

Moreover, we performed ddPCR testing *IDH2* R140Q mutation on 11 AML samples, 7 healthy donor samples, 2 commercial reference samples, 1 synthetic reference material and the wildtype sample as shown in Figures S10 to S12 and Table S7. Because AML sample #20 and healthy donor sample #7 were called positive once or twice in total nine reactions with VAF around 0.1% in ddPCR, we hypothesized that this could result from the false positive of ddPCR reactions around its LoD (Figures S11and S12). As-BDA did not report any false positive out of the nine reactions, indicating the better performance of As-BDA in specificity than that of ddPCR. From the comparative results, we concluded that As-BDA and ddPCR show 100% concordance in variant detection as shown in Table S11.

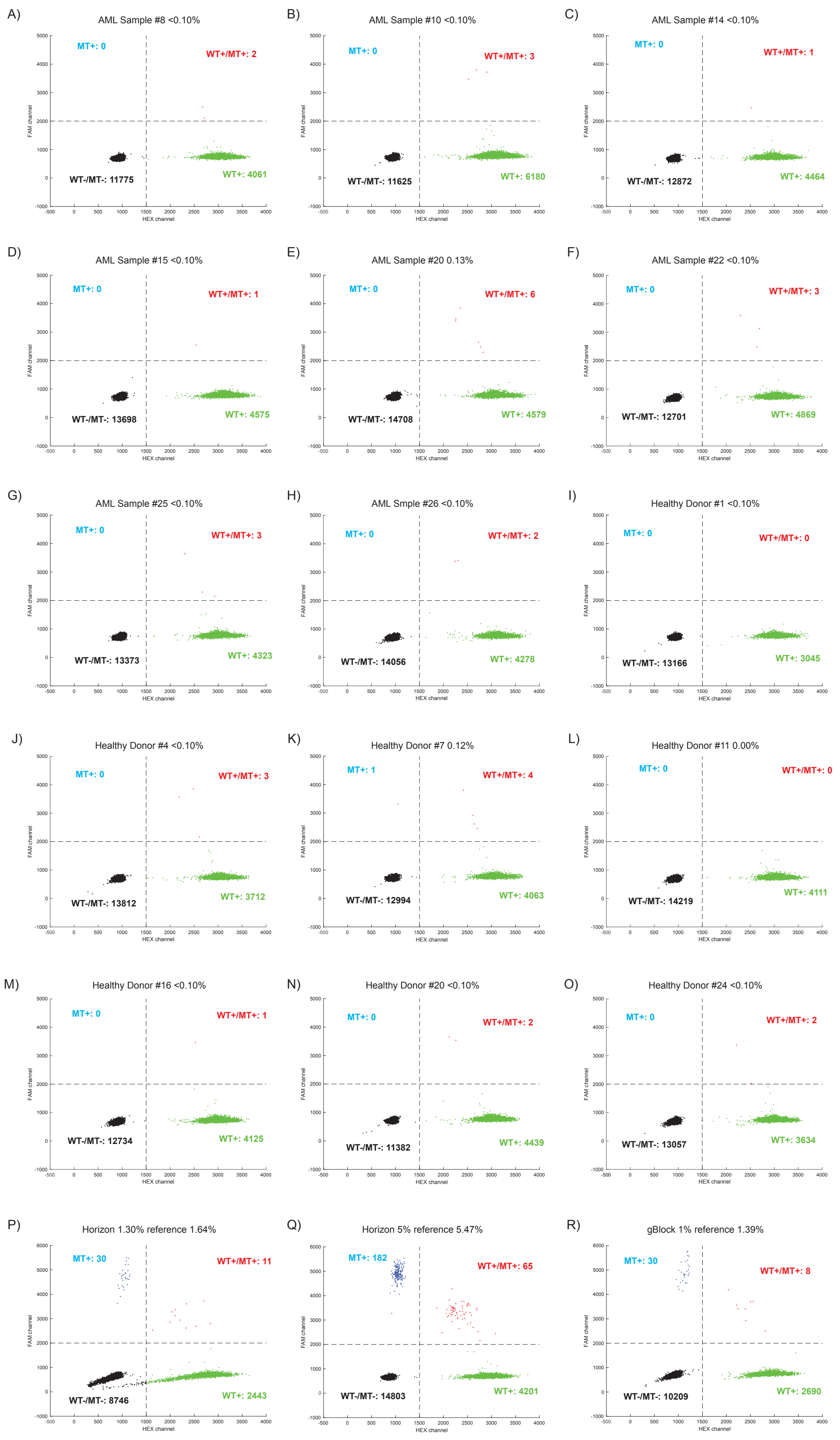


**Figure S10. ddPCR results in *IDH2* R140Q mutation.**

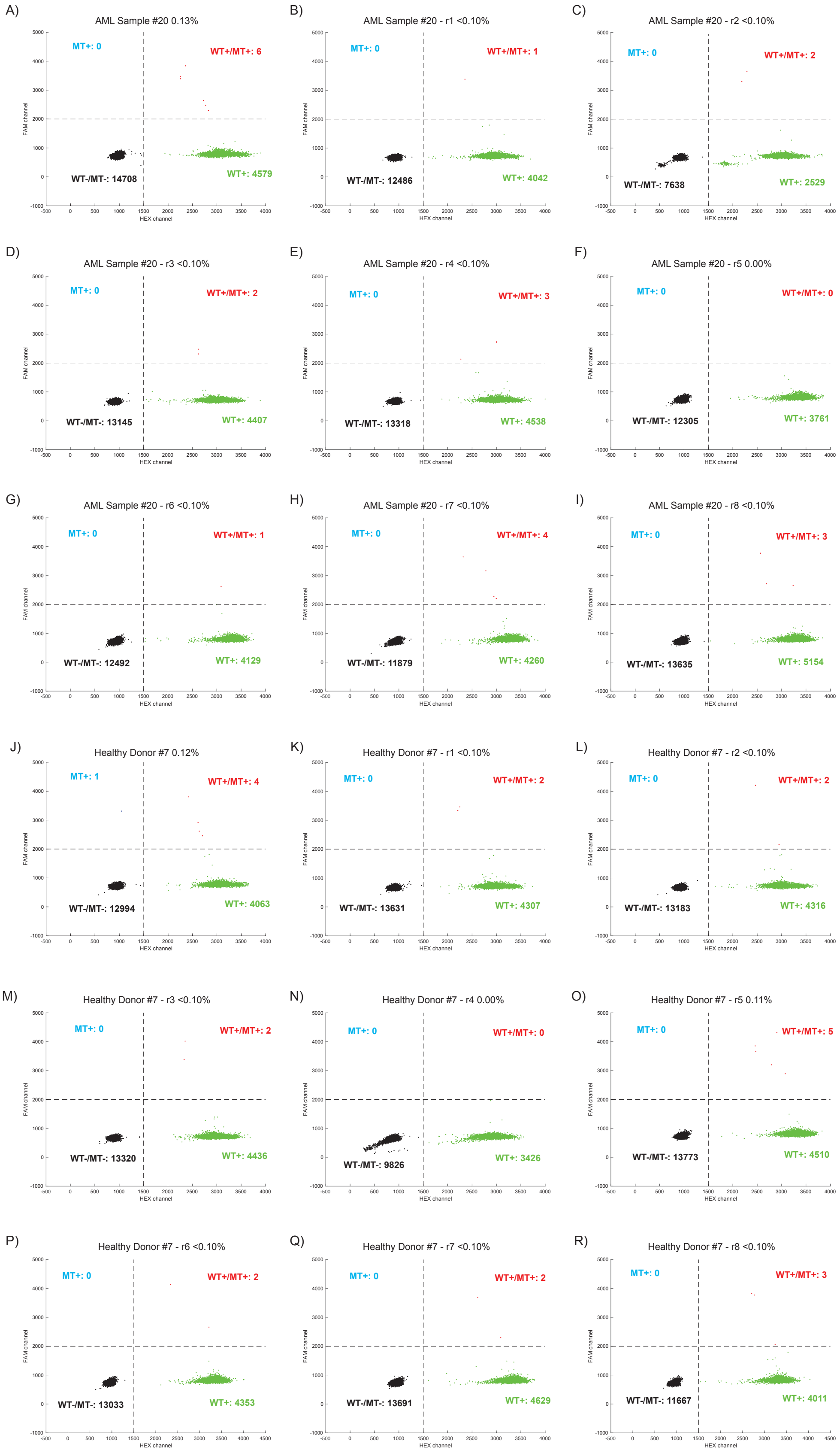
A)
AML Sample #20 0.13%
MT+: 0
WT+/MT+: 6
WT-/MT-: 14708
WT+: 4579
B)
AML Sample #20 - r1 <0.10%
MT+: 0
WT+/MT+: 1
WT-/MT-: 12486
WT+: 4042
C)
AML Sample #20 - r2 <0.10%
MT+: 0
WT+/MT+: 2
WT-/MT-: 7638
WT+: 2529
D)
AML Sample #20 - r3 <0.10%
MT+: 0
WT+/MT+: 2
WT-/MT-: 13145
WT+: 4407
E)
AML Sample #20 - r4 <0.10%
MT+: 0
WT+/MT+: 3
WT-/MT-: 13318
WT+: 4538
F)
AML Sample #20 - r5 0.00%
MT+: 0
WT+/MT+: 0
WT-/MT-: 12305
WT+: 3761
G)
AML Sample #20 - r6 <0.10%
MT+: 0
WT+/MT+: 1
WT-/MT-: 12492
WT+: 4129
H)
AML Sample #20 - r7 <0.10%
MT+: 0
WT+/MT+: 4
WT-/MT-: 11879
WT+: 4260
I)
AML Sample #20 - r8 <0.10%
MT+: 0
WT+/MT+: 3
WT-/MT-: 13635
WT+: 5154
J)
Healthy Donor #7 0.12%
MT+: 1
WT+/MT+: 4
WT-/MT-: 12994
WT+: 4063
K)
Healthy Donor #7 - r1 <0.10%
MT+: 0
WT+/MT+: 2
WT-/MT-: 13631
WT+: 4307
L)
Healthy Donor #7 - r2 <0.10%
MT+: 0
WT+/MT+: 2
WT-/MT-: 13183
WT+: 4316
M)
Healthy Donor #7 - r3 <0.10%
MT+: 0
WT+/MT+: 2
WT-/MT-: 13320
WT+: 4436
N)
Healthy Donor #7 - r4 0.00%
MT+: 0
WT+/MT+: 0
WT-/MT-: 9826
WT+: 3426
O)
Healthy Donor #7 - r5 0.11%
MT+: 0
WT+/MT+: 5
WT-/MT-: 13773
WT+: 4510
P)
Healthy Donor #7 - r6 <0.10%
MT+: 0
WT+/MT+: 2
WT-/MT-: 13033
WT+: 4353
Q)
Healthy Donor #7 - r7 <0.10%
MT+: 0
WT+/MT+: 2
WT-/MT-: 13691
WT+: 4629
R)
Healthy Donor #7 - r8 <0.10%
MT+: 0
WT+/MT+: 3
WT-/MT-: 11667
WT+: 4011
FAM channel
HEX channel

**Figure S11. ddPCR repeats of Sample #20 and Healthy donor #7 in *IDH2* R140Q mutation.** To confirm our hypothesis, we repeated the AML sample #20 and Healthy donor sample #7 each for 8 times more.

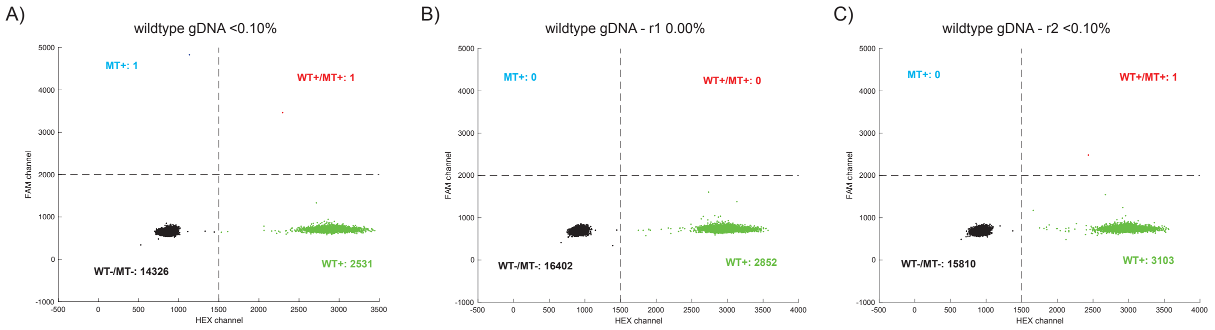


**Figure S12. ddPCR repeats of wildtype genomic DNA samples in *IDH2* R140Q mutation.** To confirm our hyphothesis, we repeated the AML sample #20 and Healthy donor sample #7 each for 8 times.

| ddPCR | Var Counts | WT Counts | VAF (%) |
|---|---|---|---|
| AML #3 | 51 | 3277 | 1.53% |
| AML #8 | 2 | 4063 | 0.05% |
| AML #10 | 3 | 6183 | 0.05% |
| AML #14 | 1 | 4465 | 0.02% |
| AML #15 | 1 | 4576 | 0.02% |
| AML #16 | 2154 | 2532 | 45.97% |
| AML #19 | 244 | 3800 | 6.03% |
| AML #20 | 6 | 4585 | 0.13% |
| AML #20 - r1 | 1 | 4042 | 0.02% |
| AML #20 - r2 | 2 | 2529 | 0.08% |
| AML #20 - r3 | 2 | 4407 | 0.05% |
| AML #20 - r4 | 3 | 4538 | 0.07% |
| AML #20 - r5 | 0 | 3760 | 0.00% |
| AML #20 - r6 | 1 | 4129 | 0.02% |
| AML #20 - r7 | 4 | 4260 | 0.09% |
| AML #20 - r8 | 3 | 5154 | 0.06% |
| AML #22 | 3 | 4869 | 0.06% |
| AML #25 | 3 | 4325 | 0.07% |
| AML #26 | 2 | 4278 | 0.05% |
| Healthy Donor #1 | 0 | 3045 | 0.00% |
| Healthy Donor #4 | 3 | 3712 | 0.08% |

| | | | |
|---|---|---|---|
| Healthy Donor #7 | 5 | 4067 | 0.12% |
| Healthy Donor #7 - r1 | 2 | 4307 | 0.05% |
| Healthy Donor #7 - r2 | 2 | 4316 | 0.05% |
| Healthy Donor #7 - r3 | 2 | 4436 | 0.05% |
| Healthy Donor #7 - r4 | 0 | 3426 | 0.00% |
| Healthy Donor #7 - r5 | 5 | 4510 | 0.11% |
| Healthy Donor #7 - r6 | 2 | 4353 | 0.05% |
| Healthy Donor #7 - r7 | 2 | 4629 | 0.04% |
| Healthy Donor #7 - r8 | 3 | 4011 | 0.07% |
| Healthy Donor #11 | 0 | 4111 | 0.00% |
| Healthy Donor #16 | 1 | 4125 | 0.02% |
| Healthy Donor #20 | 2 | 4441 | 0.05% |
| Healthy Donor #24 | 2 | 3634 | 0.06% |
| WT gDNA NA18562 | 2 | 2532 | 0.08% |
| WT gDNA NA18562 - r1 | 0 | 2852 | 0.00% |
| WT gDNA NA18562 - r2 | 1 | 3103 | 0.03% |
| 1% HD734 | 41 | 2454 | 1.64% |
| 5% HD729 | 247 | 4266 | 5.47% |
| 1% VAF ref gBlock | 38 | 2698 | 1.39% |

**Table S7. Summary of ddPCR results in R140Q mutation.** The ddPCR was analyzed using QuantaSoft software in absolute quantification (ABS) mode. Var count is from FAM channel, wildtype is from HEX channel. Double positive droplets would be counted in both var count and wt count.

| As-BDA Tube 1 - 15 ng | Texas Red Ct value | VAF (%) | Cy5 Ct value | VAF (%) | HEX Ct value | VAF (%) | Quasar Ct value | VAF (%) |
|---|---|---|---|---|---|---|---|---|
| AML #1 | Inf | 0.00 | Inf | 0.00 | Inf | 0.00 | 39.3 | 0.02 |
| AML #2 | Inf | 0.00 | Inf | 0.00 | Inf | 0.00 | 39.2 + Inf | 0.02 |
| AML #3 | Inf | 0.00 | Inf | 0.00 | Inf | 0.00 | 32.4 | 1.39 |
| AML #4 | Inf | 0.00 | 35.8 + Inf | 0.03 | Inf | 0.00 | Inf | 0.00 |
| AML #5 | Inf | 0.00 | Inf | 0.00 | Inf | 0.00 | Inf | 0.00 |
| AML #6 | Inf | 0.00 | 36.7 + Inf | 0.02 | Inf | 0.00 | Inf | 0.00 |
| AML #7 | Inf | 0.00 | 38.5 + Inf | 0.00 | Inf | 0.00 | Inf | 0.00 |
| AML #8 | Inf | 0.00 | 38.8 + Inf | 0.00 | Inf | 0.00 | Inf | 0.00 |
| AML #9 | Inf | 0.00 | Inf | 0.00 | Inf | 0.00 | Inf | 0.00 |
| AML #10 | Inf | 0.00 | Inf | 0.00 | Inf | 0.00 | Inf | 0.00 |

| | | | | | | | | |
|---|---|---|---|---|---|---|---|---|
| AML #11 | Inf | 0.00 | Inf | 0.00 | Inf | 0.00 | Inf | 0.00 |
| AML #12 | Inf | 0.00 | Inf | 0.00 | Inf | 0.00 | 41.4 + Inf | 0.00 |
| AML #13 | Inf | 0.00 | 36.5 | 0.02 | Inf | 0.00 | 36.6 + Inf | 0.09 |
| AML #14 | Inf | 0.00 | 35.5 + Inf | 0.04 | Inf | 0.00 | Inf | 0.00 |
| AML #15 | Inf | 0.00 | 37.3 + Inf | 0.01 | Inf | 0.00 | 38.1 | 0.03 |
| AML #16 | Inf | 0.00 | Inf | 0.00 | Inf | 0.00 | 26.9 | 51.22 |
| AML #17 | Inf | 0.00 | 41.3 + Inf | 0.00 | Inf | 0.00 | Inf | 0.00 |
| AML #18 | Inf | 0.00 | Inf | 0.00 | Inf | 0.00 | Inf | 0.00 |
| AML #19 | Inf | 0.00 | Inf | 0.00 | Inf | 0.00 | 30.5 | 4.83 |
| AML #20 | Inf | 0.00 | Inf | 0.00 | Inf | 0.00 | 38.7 + Inf | 0.02 |
| AML #21 | Inf | 0.00 | Inf | 0.00 | Inf | 0.00 | 39.1 + Inf | 0.02 |
| AML #22 | Inf | 0.00 | 36.6 + Inf | 0.02 | Inf | 0.00 | Inf | 0.00 |
| AML #23 | Inf | 0.00 | 35.8 + Inf | 0.03 | Inf | 0.00 | Inf | 0.00 |
| AML #24 | Inf | 0.00 | Inf | 0.00 | Inf | 0.00 | Inf | 0.00 |
| AML #25 | Inf | 0.00 | 39.4 + Inf | 0.00 | Inf | 0.00 | 38.8 + Inf | 0.02 |
| AML #26 | Inf | 0.00 | Inf | 0.00 | Inf | 0.00 | 38.1 + Inf | 0.03 |
| AML #27 | Inf | 0.00 | Inf | 0.00 | Inf | 0.00 | 36.5 + Inf | 0.09 |
| 1% HD734 | Inf | 0.00 | Inf | 0.00 | Inf | 0.00 | 32.1 | 1.69 |
| 5% HD729 | Inf | 0.00 | Inf | 0.00 | Inf | 0.00 | 30.3 | 5.50 |
| WT gDNA NA18562 | Inf | 0.00 | Inf | 0.00 | Inf | 0.00 | Inf | 0.00 |

**Table S8. Summary of Ct value and called VAF based on calibration curve of AML clinical samples, reference samples in R140 mutations.**

| As-BDA Tube 1 - 15 ng | Texas Red Ct value | VAF (%) | Cy5 Ct value | VAF (%) | HEX Ct value | VAF (%) | Quasar Ct value | VAF (%) |
|---|---|---|---|---|---|---|---|---|
| Healthy Donor #1 | Inf | 0.00 | Inf | 0.00 | Inf | 0.00 | 40.4 + Inf | 0.01 |
| Healthy Donor #2 | Inf | 0.00 | Inf | 0.00 | Inf | 0.00 | 39.8 + Inf | 0.01 |
| Healthy Donor #3 | Inf | 0.00 | Inf | 0.00 | Inf | 0.00 | 37.8 + Inf | 0.04 |
| Healthy Donor #4 | Inf | 0.00 | 38.1 + Inf | 0.01 | Inf | 0.00 | Inf | 0.00 |
| Healthy Donor #5 | Inf | 0.00 | Inf | 0.00 | Inf | 0.00 | Inf | 0.00 |
| Healthy Donor #6 | Inf | 0.00 | 40.4 + Inf | 0.00 | Inf | 0.00 | Inf | 0.00 |
| Healthy Donor #7 | Inf | 0.00 | Inf + 37.4 | 0.01 | Inf | 0.00 | 39.1 + Inf | 0.02 |
| Healthy Donor #8 | Inf | 0.00 | Inf | 0.00 | Inf | 0.00 | 38.4 + Inf | 0.03 |
| Healthy Donor #9 | Inf | 0.00 | Inf | 0.00 | Inf | 0.00 | 39.5 + Inf | 0.01 |

| | | | | | | | | |
|---|---|---|---|---|---|---|---|---|
| Healthy Donor #10 | Inf | 0.00 | 37.7 | 0.01 | Inf | 0.00 | 37.9 + Inf | 0.04 |
| Healthy Donor #11 | Inf | 0.00 | Inf | 0.00 | Inf | 0.00 | 39.0 + Inf | 0.02 |
| Healthy Donor #12 | Inf | 0.00 | 36.4 + Inf | 0.02 | Inf | 0.00 | Inf | 0.00 |
| Healthy Donor #13 | Inf | 0.00 | 39.1 + Inf | 0.00 | Inf | 0.00 | Inf | 0.00 |
| Healthy Donor #14 | Inf | 0.00 | 38.1 + Inf | 0.01 | Inf | 0.00 | 38.1 + Inf | 0.03 |
| Healthy Donor #15 | Inf | 0.00 | 37.6 + Inf | 0.01 | Inf | 0.00 | Inf | 0.00 |
| Healthy Donor #16 | Inf | 0.00 | 37.2 + Inf | 0.01 | Inf | 0.00 | Inf | 0.00 |
| Healthy Donor #17 | Inf | 0.00 | Inf | 0.00 | Inf | 0.00 | 40.6 + Inf | 0.01 |
| Healthy Donor #18 | 36.3 + Inf | 0.03 | 37.0 + Inf | 0.01 | Inf | 0.00 | Inf | 0.00 |
| Healthy Donor #19 | Inf | 0.00 | 37.3 + Inf | 0.01 | Inf | 0.00 | Inf | 0.00 |
| Healthy Donor #20 | Inf | 0.00 | 37 + Inf | 0.01 | Inf | 0.00 | Inf | 0.00 |
| Healthy Donor #21 | Inf | 0.00 | 38 + Inf | 0.01 | Inf | 0.00 | Inf | 0.00 |
| Healthy Donor #22 | Inf | 0.00 | 38.1 + Inf | 0.01 | Inf | 0.00 | Inf | 0.00 |
| Healthy Donor #23 | Inf | 0.00 | 37.3 + Inf | 0.01 | Inf | 0.00 | 37.4 + Inf | 0.05 |
| Healthy Donor #24 | Inf | 0.00 | Inf | 0.00 | Inf | 0.00 | Inf | 0.00 |

**Table S9. Summary of Ct value and called VAF based on calibration curve of healthy donor samples, reference samples in R140 mutations.** If a duplicate shows different amplification curves, then both Ct values would be collected and presented. For example, the reaction of healthy donor #18 had only one amplification curve out of a duplicate with Ct value is approximately 36.3 in Texas Red channel, then the Ct values presented are 36.3 + Inf. The smaller Ct value would be used for VAF calling based on the regression equation obtained from synthetic reference materials.

| As-BDA - 15 ng<br>Tube 1 - Quasar | Ct value | VAF (%) |
|---|---|---|
| AML #20 | 38.7 + Inf | 0.02 |
| AML #20 - r1 | Inf | 0.00 |
| AML #20 - r2 | Inf | 0.00 |
| AML #20 - r3 | Inf | 0.00 |
| AML #20 - r4 | Inf | 0.00 |
| AML #20 - r5 | Inf | 0.00 |
| AML #20 - r6 | Inf | 0.00 |
| AML #20 - r7 | Inf | 0.00 |
| AML #20 - r8 | Inf | 0.00 |
| Healthy Donor #7 | 39.1 + Inf | 0.02 |
| Healthy Donor #7 - r1 | 38.1 + Inf | 0.03 |

| | | |
|---|---|---|
| Healthy Donor #7 - r2 | Inf | 0.00 |
| Healthy Donor #7 - r3 | Inf | 0.00 |
| Healthy Donor #7 - r4 | Inf | 0.00 |
| Healthy Donor #7 - r5 | Inf | 0.00 |
| Healthy Donor #7 - r6 | 39.2 + Inf | 0.02 |
| Healthy Donor #7 - r7 | Inf | 0.00 |
| Healthy Donor #7 - r8 | Inf | 0.00 |

**Table S10. Summary of Ct value and called VAF of repeated AML sample #20 and healthy donor sample #7 in R140Q mutation.**

| Sample ID | ddPCR VAF | As-BDA called VAF |
|---|---|---|
| AML #3 | 1.53% | 1.39% |
| AML #8 | 0.05% | 0.00% |
| AML #10 | 0.05% | 0.00% |
| AML #14 | 0.02% | 0.00% |
| AML #15 | 0.02% | 0.03% |
| AML #16 | 45.97% | 49.89% |
| AML #19 | 6.03% | 4.83% |
| AML #20 | 0.06% | 0.02% |
| AML #22 | 0.06% | 0.00% |
| AML #25 | 0.07% | 0.02% |
| AML #26 | 0.05% | 0.03% |
| 5% HD729 | 5.47% | 5.50% |
| 1% HD734 | 1.64% | 1.69% |
| WT gDNA NA18562 | 0.03% | 0.00% |
| 1% VAF ref gBlock | 1.39% | / |
| Healthy Donor #1 | 0.00% | 0.01% |
| Healthy Donor #4 | 0.08% | 0.00% |
| Healthy Donor #7 | 0.05% | 0.02% |
| Healthy Donor #11 | 0.00% | 0.02% |
| Healthy Donor #16 | 0.02% | 0.00% |
| Healthy Donor #20 | 0.05% | 0.00% |
| Healthy Donor #24 | 0.06% | 0.00% |

**Table S11. Clinical sample summary of comparative analytical results in *IDH2* R140Q mutation.** Sample results are shown for As-BDA and ddPCR.

NGS was performed on 3 AML clinical samples, 2 commercial reference samples, 2 synthetic reference materials and the wildtype sample targeting R140 region (Table S12 to S15).

| NGS | Var Reads | WT Reads | VAF (%) |
|---|---|---|---|
| WT gDNA NA18562 | 73 | 654555 | 0.01% |
| WT gDNA NA18562 | 101 | 801480 | 0.01% |
| 10% VAF ref gBlock | 2913 | 29305 | 9.04% |
| 1% VAF ref gBlock | 2842 | 179497 | 1.56% |

**Table S12. Summary of VAF from NGS in R140 418 C>G mutation.**

| NGS | Var Reads | WT Reads | VAF (%) |
|---|---|---|---|
| WT gDNA NA18562 | 766 | 654555 | 0.12% |
| WT gDNA NA18562 | 1022 | 801480 | 0.13% |
| 10% VAF ref gBlock | 1352 | 9350 | 12.63% |
| 1% VAF ref gBlock | 3599 | 181660 | 1.94% |

**Table S13. Summary of VAF from NGS in R140 418 C>T mutation.**

| NGS | Var Reads | WT Reads | VAF (%) |
|---|---|---|---|
| WT gDNA NA18562 | 234 | 654555 | 0.04% |
| WT gDNA NA18562 | 402 | 801480 | 0.05% |
| 10% VAF ref gBlock | 2545 | 23941 | 9.61% |
| 1% VAF ref gBlock | 3251 | 236829 | 1.35% |

**Table S14. Summary of VAF from NGS in R140 419 G>T mutation.**

| NGS | Var Reads | WT Reads | VAF (%) |
|---|---|---|---|
| AML #3 | 12595 | 863060 | 1.44% |
| AML #16 | 392361 | 472739 | 45.35% |
| AML #19 | 42994 | 765884 | 5.32% |
| 1.3% HD734 | 10055 | 675300 | 1.47% |
| 5% HD729 | 48198 | 708262 | 6.37% |

| | | | |
|---|---|---|---|
| WT gDNA NA18562 | 817 | 801480 | 0.10% |
| WT gDNA NA18562 | 1696 | 654555 | 0.26% |
| 1% VAF ref gBlock | 3644 | 205943 | 1.74% |

**Table S15. Summary of VAF from NGS in R140 419 G>A mutation.**

The obtained Ct values and accordingly called VAF of tube 2 and 3 were presented in Table S16 to S19. For R172 region, 2 commercial reference samples, 2 synthetic reference materials and the wildtype sample were tested on NGS (Table S20 to S25). As-BDA reactions did not report any AML patients or healthy donors with R172 mutations.

| As-BDA Tube 2 - 15 ng | Cy5 Ct value | VAF (%) | Texas Red Ct value | VAF (%) | HEX Ct value | VAF (%) | Quasar Ct value |
|---|---|---|---|---|---|---|---|
| AML #1 | Inf | 0.00 | Inf | 0.00 | Inf | 0.00 | 31.7 |
| AML #2 | Inf | 0.00 | Inf | 0.00 | Inf | 0.00 | 31.7 |
| AML #3 | Inf | 0.00 | Inf | 0.00 | Inf | 0.00 | 31.7 |
| AML #4 | Inf | 0.00 | Inf | 0.00 | Inf | 0.00 | 31.3 |
| AML #5 | Inf | 0.00 | Inf | 0.00 | Inf | 0.00 | 31.9 |
| AML #6 | Inf | 0.00 | Inf | 0.00 | Inf | 0.00 | 32.4 |
| AML #7 | Inf | 0.00 | Inf | 0.00 | Inf | 0.00 | 32.1 |
| AML #8 | Inf | 0.00 | Inf | 0.00 | Inf | 0.00 | 32.1 |
| AML #9 | Inf | 0.00 | Inf | 0.00 | Inf | 0.00 | 32.3 |
| AML #10 | Inf | 0.00 | Inf | 0.00 | Inf | 0.00 | 31.2 |
| AML #11 | Inf | 0.00 | Inf | 0.00 | Inf | 0.00 | 32.6 |
| AML #12 | Inf | 0.00 | Inf | 0.00 | Inf | 0.00 | 32.0 |
| AML #13 | Inf | 0.00 | Inf | 0.00 | Inf | 0.00 | 31.4 |
| AML #14 | Inf | 0.00 | Inf | 0.00 | Inf | 0.00 | 31.8 |
| AML #15 | Inf | 0.00 | Inf | 0.00 | Inf | 0.00 | 32.1 |
| AML #16 | Inf | 0.00 | Inf | 0.00 | Inf | 0.00 | 31.9 |
| AML #17 | Inf | 0.00 | Inf | 0.00 | Inf | 0.00 | 31.6 |
| AML #18 | 38.8 + Inf | 0.00 | Inf | 0.00 | Inf | 0.00 | 31.8 |
| AML #19 | Inf | 0.00 | Inf | 0.00 | Inf | 0.00 | 32.1 |
| AML #20 | Inf | 0.00 | Inf | 0.00 | Inf | 0.00 | 32.1 |
| AML #21 | Inf | 0.00 | Inf | 0.00 | Inf | 0.00 | 32.0 |
| AML #22 | Inf | 0.00 | Inf | 0.00 | Inf | 0.00 | 31.5 |

| | | | | | | | |
|---|---|---|---|---|---|---|---|
| AML #23 | Inf | 0.00 | Inf | 0.00 | Inf | 0.00 | 32.1 |
| AML #24 | Inf | 0.00 | Inf | 0.00 | Inf | 0.00 | 31.3 |
| AML #25 | Inf | 0.00 | Inf | 0.00 | Inf | 0.00 | 31.9 |
| AML #26 | Inf | 0.00 | Inf | 0.00 | Inf | 0.00 | 32.1 |
| AML #27 | Inf | 0.00 | Inf | 0.00 | Inf | 0.00 | 31.7 |
| 1% HD734 | Inf | 0.00 | Inf | 0.00 | Inf | 0.00 | 32.6 |
| 1% HD829 | Inf | 0.00 | Inf | 0.00 | Inf | 0.00 | 32.8 |
| 5% HD729 | Inf | 0.00 | Inf | 0.00 | Inf | 0.00 | 32.7 |
| WT gDNA NA18562 | Inf | 0.00 | Inf | 0.00 | Inf | 0.00 | 32.1 |

**Table S16. Summary of Ct value and called VAF based on calibration curve of AML clinical samples, reference samples in R172 mutations in tube 2.**

| As-BDA Tube 2 - 15 ng | Cy5 Ct value | VAF (%) | Texas Red Ct value | VAF (%) | HEX Ct value | VAF (%) | Quasar Ct value |
|---|---|---|---|---|---|---|---|
| Healthy Donor #1 | Inf | 0.00 | Inf | 0.00 | Inf | 0.00 | 32.4 |
| Healthy Donor #2 | Inf | 0.00 | Inf | 0.00 | Inf | 0.00 | 32.1 |
| Healthy Donor #3 | Inf | 0.00 | Inf | 0.00 | 43.5 + Inf | 0.00 | 32.6 |
| Healthy Donor #4 | Inf | 0.00 | Inf | 0.00 | Inf | 0.00 | 32.6 |
| Healthy Donor #5 | Inf | 0.00 | Inf | 0.00 | Inf | 0.00 | 32.4 |
| Healthy Donor #6 | Inf | 0.00 | Inf | 0.00 | Inf | 0.00 | 32.5 |
| Healthy Donor #7 | Inf | 0.00 | Inf | 0.00 | Inf | 0.00 | 31.0 |
| Healthy Donor #8 | Inf | 0.00 | Inf | 0.00 | Inf | 0.00 | 31.7 |
| Healthy Donor #9 | Inf | 0.00 | Inf | 0.00 | Inf | 0.00 | 32.1 |
| Healthy Donor #10 | Inf | 0.00 | Inf | 0.00 | Inf | 0.00 | 32.3 |
| Healthy Donor #11 | Inf | 0.00 | Inf | 0.00 | Inf | 0.00 | 32.3 |
| Healthy Donor #12 | Inf | 0.00 | Inf | 0.00 | Inf | 0.00 | 32.1 |
| Healthy Donor #13 | Inf | 0.00 | Inf | 0.00 | Inf | 0.00 | 32.3 |
| Healthy Donor #14 | Inf | 0.00 | Inf | 0.00 | Inf | 0.00 | 32.2 |
| Healthy Donor #15 | Inf | 0.00 | Inf | 0.00 | Inf | 0.00 | 32.0 |
| Healthy Donor #16 | Inf | 0.00 | Inf | 0.00 | Inf | 0.00 | 31.7 |
| Healthy Donor #17 | Inf | 0.00 | Inf | 0.00 | Inf | 0.00 | 32.1 |
| Healthy Donor #18 | Inf | 0.00 | Inf | 0.00 | Inf | 0.00 | 32.0 |
| Healthy Donor #19 | Inf | 0.00 | Inf | 0.00 | 57.8 + Inf | 0.00 | 31.7 |
| Healthy Donor #20 | Inf | 0.00 | Inf | 0.00 | Inf | 0.00 | 31.7 |

| | | | | | | | |
|---|---|---|---|---|---|---|---|
| Healthy Donor #21 | Inf | 0.00 | Inf | 0.00 | Inf | 0.00 | 31.6 |
| Healthy Donor #22 | Inf | 0.00 | Inf | 0.00 | Inf | 0.00 | 31.9 |
| Healthy Donor #23 | Inf | 0.00 | Inf | 0.00 | Inf | 0.00 | 31.7 |
| Healthy Donor #24 | Inf | 0.00 | Inf | 0.00 | Inf | 0.00 | 32.2 |

**Table S17. Summary of Ct value and called VAF based on calibration curve of healthy donor clinical samples, reference samples in R172 mutations in tube 2.**

| As-BDA Tube 3 - 15 ng | Cy5 Ct value | VAF (%) | Texas Red Ct value | VAF (%) | HEX Ct value | VAF (%) | Quasar Ct value |
|---|---|---|---|---|---|---|---|
| AML #1 | Inf | 0.00 | Inf | 0.00 | Inf | 0.00 | 31.9 |
| AML #2 | Inf | 0.00 | Inf | 0.00 | Inf | 0.00 | 31.9 |
| AML #3 | Inf | 0.00 | Inf | 0.00 | Inf | 0.00 | 31.9 |
| AML #4 | Inf | 0.00 | Inf | 0.00 | Inf | 0.00 | 31.2 |
| AML #5 | Inf | 0.00 | Inf | 0.00 | Inf | 0.00 | 31.8 |
| AML #6 | Inf | 0.00 | Inf | 0.00 | Inf | 0.00 | 32.4 |
| AML #7 | Inf | 0.00 | Inf | 0.00 | Inf | 0.00 | 32.1 |
| AML #8 | Inf | 0.00 | Inf | 0.00 | Inf | 0.00 | 32.0 |
| AML #9 | Inf | 0.00 | Inf | 0.00 | Inf | 0.00 | 32.0 |
| AML #10 | Inf | 0.00 | Inf | 0.00 | Inf | 0.00 | 31.1 |
| AML #11 | Inf | 0.00 | Inf | 0.00 | Inf | 0.00 | 32.6 |
| AML #12 | Inf | 0.00 | Inf | 0.00 | Inf | 0.00 | 32.1 |
| AML #13 | Inf | 0.00 | Inf | 0.00 | Inf | 0.00 | 31.0 |
| AML #14 | Inf | 0.00 | Inf | 0.00 | Inf | 0.00 | 32.1 |
| AML #15 | Inf | 0.00 | Inf | 0.00 | Inf | 0.00 | 32.0 |
| AML #16 | Inf | 0.00 | Inf | 0.00 | Inf | 0.00 | 31.9 |
| AML #17 | Inf | 0.00 | Inf | 0.00 | Inf | 0.00 | 31.6 |
| AML #18 | Inf | 0.00 | Inf | 0.00 | Inf | 0.00 | 32.0 |
| AML #19 | Inf | 0.00 | Inf | 0.00 | Inf | 0.00 | 31.9 |
| AML #20 | Inf | 0.00 | Inf | 0.00 | Inf | 0.00 | 32.2 |
| AML #21 | Inf | 0.00 | Inf | 0.00 | Inf | 0.00 | 32.0 |
| AML #22 | Inf | 0.00 | Inf | 0.00 | Inf | 0.00 | 31.6 |
| AML #23 | Inf | 0.00 | Inf | 0.00 | Inf | 0.00 | 32.0 |
| AML #24 | Inf | 0.00 | Inf | 0.00 | Inf | 0.00 | 31.8 |
| AML #25 | Inf | 0.00 | Inf | 0.00 | Inf | 0.00 | 32.2 |
| AML #26 | Inf | 0.00 | Inf | 0.00 | 51.1 + Inf | 0.00 | 32.2 |

| | | | | | | | |
|---|---|---|---|---|---|---|---|
| AML #27 | Inf | 0.00 | Inf | 0.00 | Inf | 0.00 | 31.9 |
| 1% HD734 | Inf | 0.00 | Inf | 0.00 | Inf | 0.00 | 32.6 |
| 1% HD829 | Inf | 0.00 | 33.5 | 1.25 | Inf | 0.00 | 32.5 |
| 5% HD729 | Inf | 0.00 | 30.9 | 5.41 | Inf | 0.00 | 32.4 |
| WT gDNA NA18562 | Inf | 0.00 | Inf | 0.00 | Inf | 0.00 | 31.9 |

**Table S18. Summary of Ct value and called VAF based on calibration curve of AML clinical samples, reference samples in R172 mutations in tube 3.**

| As-BDA Tube 3 - 15 ng | Cy5 Ct value | VAF (%) | Texas Red Ct value | VAF (%) | HEX Ct value | VAF (%) | Quasar Ct value |
|---|---|---|---|---|---|---|---|
| Healthy Donor #1 | Inf | 0.00 | Inf | 0.00 | Inf | 0.00 | 32.3 |
| Healthy Donor #2 | Inf | 0.00 | Inf | 0.00 | Inf | 0.00 | 31.9 |
| Healthy Donor #3 | Inf | 0.00 | Inf | 0.00 | Inf | 0.00 | 32.4 |
| Healthy Donor #4 | Inf | 0.00 | Inf | 0.00 | Inf | 0.00 | 32.3 |
| Healthy Donor #5 | Inf | 0.00 | Inf | 0.00 | Inf | 0.00 | 32.3 |
| Healthy Donor #6 | Inf | 0.00 | Inf | 0.00 | Inf | 0.00 | 32.7 |
| Healthy Donor #7 | Inf | 0.00 | Inf | 0.00 | Inf | 0.00 | 31.8 |
| Healthy Donor #8 | Inf | 0.00 | Inf | 0.00 | 49.3 + Inf | 0.00 | 31.5 |
| Healthy Donor #9 | Inf | 0.00 | Inf | 0.00 | Inf | 0.00 | 32.2 |
| Healthy Donor #10 | Inf | 0.00 | Inf | 0.00 | Inf | 0.00 | 32.3 |
| Healthy Donor #11 | Inf | 0.00 | Inf | 0.00 | Inf | 0.00 | 32.4 |
| Healthy Donor #12 | Inf | 0.00 | Inf | 0.00 | Inf | 0.00 | 32.3 |
| Healthy Donor #13 | Inf | 0.00 | Inf | 0.00 | Inf | 0.00 | 32.7 |
| Healthy Donor #14 | Inf | 0.00 | Inf | 0.00 | Inf | 0.00 | 32.5 |
| Healthy Donor #15 | Inf | 0.00 | Inf | 0.00 | Inf | 0.00 | 31.9 |
| Healthy Donor #16 | Inf | 0.00 | Inf | 0.00 | Inf | 0.00 | 32.1 |
| Healthy Donor #17 | Inf | 0.00 | Inf | 0.00 | Inf | 0.00 | 32.1 |
| Healthy Donor #18 | Inf | 0.00 | Inf | 0.00 | Inf | 0.00 | 31.8 |
| Healthy Donor #19 | Inf | 0.00 | Inf | 0.00 | Inf | 0.00 | 31.7 |
| Healthy Donor #20 | Inf | 0.00 | Inf | 0.00 | Inf | 0.00 | 31.5 |
| Healthy Donor #21 | Inf | 0.00 | Inf | 0.00 | Inf | 0.00 | 31.5 |
| Healthy Donor #22 | Inf | 0.00 | Inf | 0.00 | Inf | 0.00 | 32.0 |
| Healthy Donor #23 | Inf | 0.00 | Inf | 0.00 | Inf | 0.00 | 31.8 |
| Healthy Donor #24 | Inf | 0.00 | Inf | 0.00 | 51.6 + Inf | 0.00 | 32.2 |

**Table S19. Summary of Ct value and called VAF based on calibration curve of healthy donor clinical samples, reference samples in R172 mutations in tube 3.**

| NGS | Var Reads | WT Reads | VAF (%) |
|---|---|---|---|
| WT gDNA NA18562 | 80 | 901487 | 0.01% |
| 10% VAF ref gBlock | 10213 | 96885 | 9.54% |
| 1% VAF ref gBlock | 3784 | 405158 | 0.93% |

**Table S20. Summary of VAF from NGS in R172 516 G>C mutation in tube 2.**

| NGS | Var Reads | WT Reads | VAF (%) |
|---|---|---|---|
| WT gDNA NA18562 | 106 | 901487 | 0.01% |
| 10% VAF ref gBlock | 6117 | 67496 | 8.31% |
| 1% VAF ref gBlock | 2984 | 408794 | 0.72% |

**Table S21. Summary of VAF from NGS in R172 515 G>T mutation in tube 2.**

| NGS | Var Reads | WT Reads | VAF (%) |
|---|---|---|---|
| WT gDNA NA18562 | 512 | 901487 | 0.06% |
| 10% VAF ref gBlock | 2615 | 25163 | 9.41% |
| 1% VAF ref gBlock | 252 | 20864 | 1.19% |

**Table S22. Summary of VAF from NGS in R172 514 A>T mutation in tube 2.**

| NGS | Var Reads | WT Reads | VAF (%) |
|---|---|---|---|
| WT gDNA NA18562 | 120 | 901487 | 0.01% |
| 10% VAF ref gBlock | 7584 | 90942 | 7.70% |
| 1% VAF ref gBlock | 267 | 25924 | 1.02% |

**Table S23. Summary of VAF from NGS in R172 516 G>T mutation in tube 3.**

| NGS | Var Reads | WT Reads | VAF (%) |
|---|---|---|---|
| 5% HD829 | 53272 | 1055183 | 4.81% |
| 1.3% HD734 | 13234 | 925099 | 1.41% |
| WT gDNA NA18562 | 859 | 901487 | 0.10% |
| 10% VAF ref gBlock | 10713 | 76765 | 12.25% |
| 1% VAF ref gBlock | 428 | 35310 | 1.20% |

**Table S24. Summary of VAF from NGS in R172 515 G>A mutation in tube 3.**

| NGS | Var Reads | WT Reads | VAF (%) |
|---|---|---|---|
| WT gDNA NA18562 | 1630 | 901487 | 0.18% |
| 5% VAF ref gBlock | 4571 | 89153 | 4.88% |
| 1% VAF ref gBlock | 3784 | 405158 | 0.93% |

**Table S25. Summary of VAF from NGS in R172 514 A>G mutation in tube 3.**

### **Section S7.** **Results of Sanger sequencing.**

We double confirmed the identity of detected variants from As-BDA in Sanger Sequencing. The three AML samples were confirmed with R140Q (419 G>A) mutation as compared with the sequence of the wildtype gDNA (Figure S13).

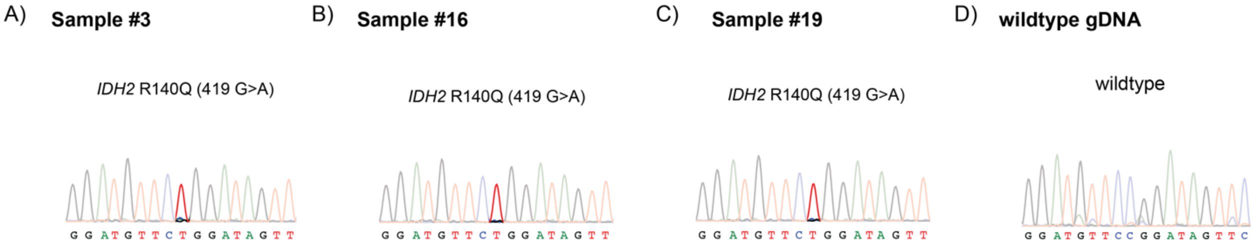


**Figure S13. Sanger Sequencing validation of 3 AML patients with R140Q mutation and wildtype sample.**

Figures S14 to S16 show the Sanger traces of synthetic reference samples post-As-BDA reactions in detecting 0.1% VAFs with 15 ng DNA input.

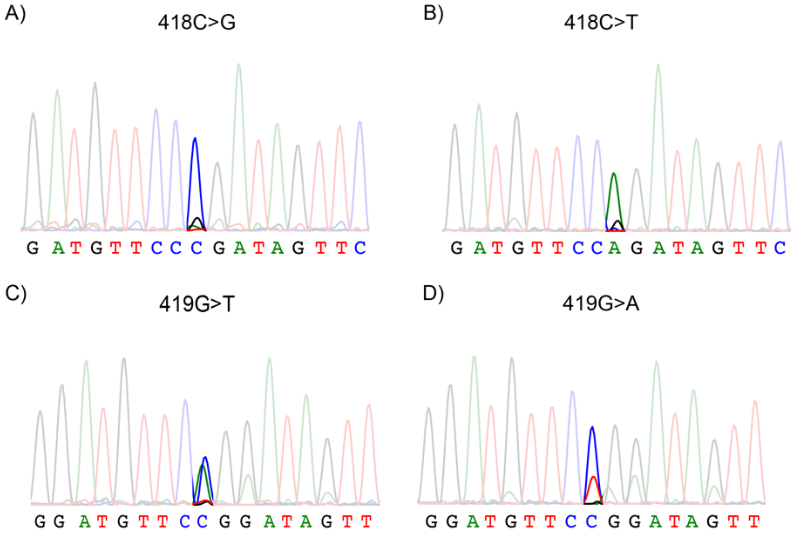


**Figure S14. Sanger Sequencing confirmation of 0.1% VAFs of tube 1.**

**(A). Sanger trace of 0.1% VAF *IDH2* 140 418C>G synthetic reference sample.** The Sanger trace shows the enrichment of G allele versus C allele because R140 primers target the minus strand.

**(B). Sanger trace of 0.1% VAF *IDH2* 140 418C>T synthetic reference sample.** The Sanger trace shows the enrichment of G allele versus A allele because R140 primers target the minus strand.

**(C). Sanger trace of 0.1% VAF *IDH2* 140 419G>T synthetic reference sample.** The Sanger trace shows the enrichment of C allele versus A allele because R140 primers target the minus strand.

**(D). Sanger trace of 0.1% VAF *IDH2* 140 419G>A synthetic reference sample.** The Sanger trace shows the enrichment of C allele versus T allele because R140 primers target the minus strand.

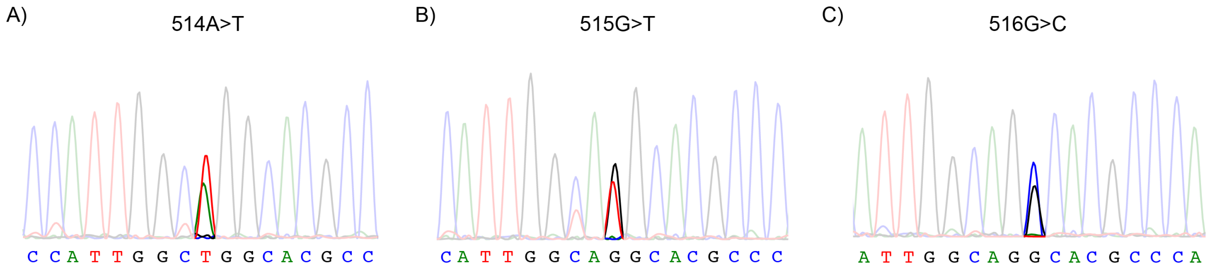


**Figure S15. Sanger Sequencing confirmation of 0.1% VAFs of tube 2.**

**(A). Sanger trace of 0.1% VAF *IDH2* 172 514A>T synthetic reference sample.**

**(B). Sanger trace of 0.1% VAF *IDH2* 172 515G>T synthetic reference sample.**

**(C). Sanger trace of 0.1% VAF *IDH2* 172 516G>C synthetic reference sample.**

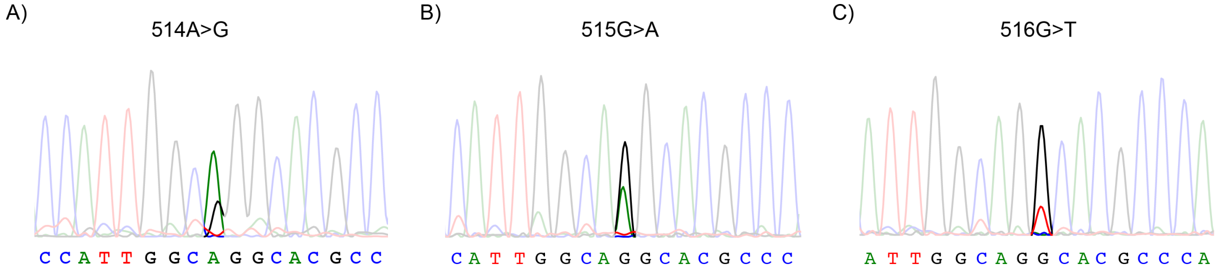


**Figure S16. Sanger Sequencing confirmation of 0.1% VAFs of tube 3.**

**(A). Sanger trace of 0.3% VAF *IDH2* 172 514A>T synthetic reference sample.**

**(B). Sanger trace of 0.1% VAF *IDH2* 172 515G>A synthetic reference sample.**

**(C). Sanger trace of 0.3% VAF *IDH2* 172 516G>T synthetic reference sample.**

However, for mutations 514 A>G and 516 G>T, we could only detect traces of variant sequences from 0.3% VAFs (Figure S16). We hypothesized that the failure of detecting 0.1% VAF 514 A>G and 0.1% VAF 516 G>T mutations in Sanger sequencing was probably because the LoD of Sanger sequencing is approximately 15 % to 20 %. The enrichment performance of the same BDA design is varied on individual mutations. The generation of the qPCR curve of 0.1% VAF 514 A>G and 0.1% VAF 516 G>T and the flat fluorescence signal of wildtype sample in Figure S6 indicated the existence of the mutation and the success of mutation detection. While, the low final fluorescence of qPCR curves also implies the inadequate number of enriched amplicons, which is likely to be below the LoD of Sanger sequencing.

We further validated our hypothesis by sequencing the qPCR product of 0.1 % VAF 514 A>G and 0.1% VAF 516 G>T (Table S26). We appended sequencing adaptor and index to the qPCR products and sequenced them in 2 x 75 bp paired-end sequencing. The NGS results confirmed the existence of enriched variants in the final library, but the variant read fraction (VRF) is only 4.54% for 514 A>G sample and 6.79%

for 516 G>T sample which are much lower than the LoD of Sanger sequencing. We further calculated the enrichment fold (EF) based on the formula:

$$\mathrm{EF} = \frac{\mathrm{VRF}\ (\mathrm{VAF}-1)}{\mathrm{VAF}\ (\mathrm{VRF}-1)}$$

The EF of As-BDA in 516 G>T is 72.8 and that in 514 A>G is around 47.5, much lower than the median of 300-fold reported before[2]. These results demonstrated our hypothesis that the As-BDA design has low enrichment effectiveness in these two mutations, making the final concentration of variant amplicons below the LoD of Sanger sequencing, leading to the failure of detection in Sanger sequencing. In this way, we could claim that our method is more sensitive and less in false negative than the previous BDA followed by Sanger sequencing[3].

| Library | WT reads | 516G>T | 514A>G | VAF |
|---|---|---|---|---|
| 0.1% VAF 516G>T | 856,433 | 62,422 | / | 6.79% |
| 0.1% VAF 514A>G | 1,241,427 | / | 58,991 | 4.54% |

**Table S26. Summary of NGS results of 0.1% VAF sample in 516 G>A and 514 A>G.**

Figure S17 shows the Sanger traces of synthetic reference samples post-As-BDA reactions in detecting 0.01% VAFs with 160 ng DNA input. Sanger trace of 0.01% VAF of mutation 516 G>C is suspicious of the same reason as described above for the failure to be detected in Sanger sequencing.

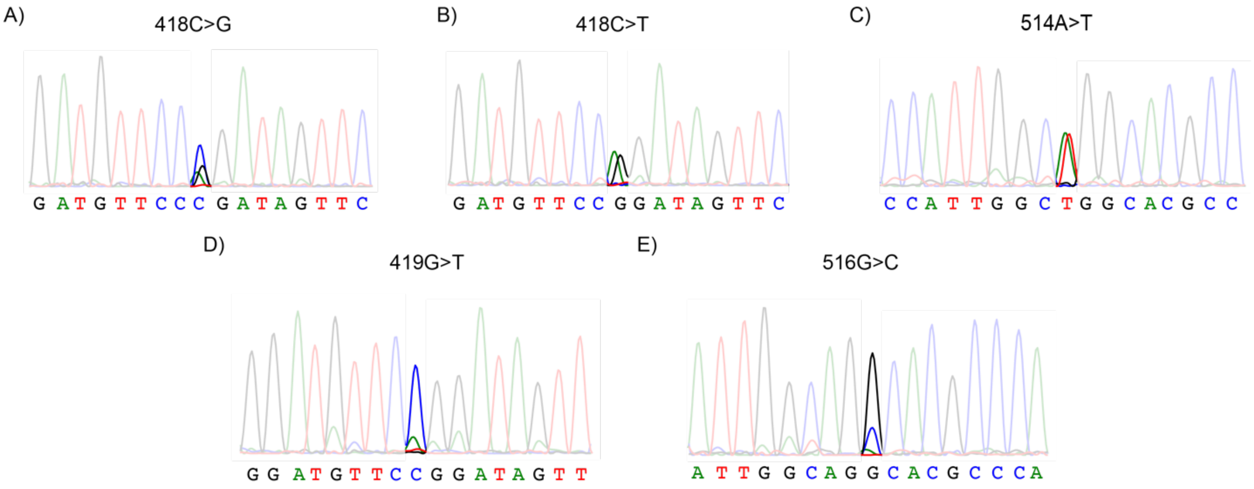


**Figure S17. Sanger Sequencing confirmation of 0.01% VAFs.**

**(A). Sanger trace of 0.01% VAF *IDH2* R140 418C>G synthetic reference sample.**

**(B). Sanger trace of 0.01% VAF *IDH2* R140 418C>T synthetic reference sample.**

**(C). Sanger trace of 0.01% VAF *IDH2* R172 514A>T synthetic reference sample.**

**(D). Sanger trace of 0.01% VAF *IDH2* R140 419G>T synthetic reference sample.**

**(E). Sanger trace of 0.03% VAF *IDH2* R172 516G>C synthetic reference sample.**

### Section S8. List of As-BDA Oligonucleotide Sequences.

The sequences of oligonucleotides and synthetic gBlocks used were attached below. The '+' symbol indicates the locked nucleic acid chemistries included in the oligo synthesis.

| Name | Sequence | Tube # |
|---|---|---|
| *IDH2*_140_FP | AGAAGATGTGGAAAAGTCCCAATG | 1 |
| *IDH2*_140_RP | GTGCCCAGGTCAGTGGAT | |
| *IDH2*_140_Blocker | TCCCAATGGAACTATCCGGAACATCC/3SpC3/ | |
| *IDH2*-140-419GT-TM | /5HEX/CTATCC+TGAACATCCTGG/3IABkFQ/ | |
| *IDH2*-140-418CT-TM | /5Cy5/CTATC+TGGAACATCCTGG/3IAbRQSp/ | |
| *IDH2*-140-418CG-TM | /56-ROXN/CTATCGGGAACATCCTGG/3IAbRQSp/ | |
| *IDH2*-140-419GA-TM | /5Cy55/ACTATCCAGAACATCCTGG/3IAbRQSp/ | |
| *IDH2*_172_FP | CTGGTCGCCATGGGCGT | 2 & 3 |
| *IDH2*_172_RP | TGAAGAAGATGTGGAAAAGTCCCA | |
| *IDH2*_172_Blocker | GGCGTGCCTGCCAATGGTGTAAAA | |
| *GAPDH*_FP3 | CAACTACATGGTGAGTGCTACATG | |
| *GAPDH*_RP3 | ATTTGCCATGGGTGGAATCATATT | |
| *GAPDH*_Blocker | GTGCTACATGGTGAGCCCCAAAGCTAAAAA | |
| *GAPDH*_TM | /5Cy55/CAGGTGGCTCCTCCCACACCAGCTT/3IAbRQSp/ | |
| *IDH2*-172-516GC-TM | /5Cy5/CGTG+GCTGCCAATGG/3IAbRQSp/ | 2 |
| *IDH2*-172-515GT-TM | /5TexRd-XN/CGTGCATGCCAATGGTG/3IAbRQSp/ | |
| *IDH2*-172-514AT-TM | /5HEX/CGTGCC+AGCCAATGG/3IABkFQ/ | |
| *IDH2*-172-516GT-TM | /5Cy5/CGTG+ACTGCCAATGGTG/3IAbRQSp/ | 3 |
| *IDH2*-172-515GA-TM | /5TexRd-XN/CGTGCTTGCCAATGGT/3IAbRQSp/ | |
| *IDH2*-172-514AG-TM | /5HEX/CGTGCC+CGCCAATGGT/3IABkFQ/ | |
| rev_*IDH2*-140-418CT | /5Cy5/CCAGGATGTTCC+AGATAGT/3IAbRQSp/ | Supple |

**Table S27. Sequence of oligonucleotides used in the As-BDA *IDH2* assay.**

| | |
|---|---|
| IDH2_418C>G | CAGACTCCAGAGCCCACACATTTGCACTCTAGACTCTACTGCCTTCCTCATGAAGAATTTTAGGACCCCCGTCTGGCTGTGTTGTTGCTTGGGGTTCAAAT<br>TCTGGTTGAAAGATGGCGGCTGCAGTGGGACCACTATTATCTCTGTCCTCACAGAGTTCAAGCTGAAGAAGATGTGGAAAAGTCCCAATGGAACTATCgGG<br>AACATCCTGGGGGGGACTGTCTTCCGGGAGCCCATCATCTGCAAAAACATCCCACGCCTAGTCCCTGGCTGGACCAAGCCCATCACCATTGGCAGGCACGC<br>CCATGGCGACCAGGTAGGCCAGGGTGGAGAGGGGATCCACTGACCTGGGCACCCCCCGACTGGAGCTCCTCGCCTAGCCATCCTCTTGTCTCTGCAGTACA<br>AGGCCACAGACTTTGTGGCAGACCGGGCCGGCACTTTCAAAATGGTCTTCACCCCAAAAGATGGCAGTGGTGTCAAGGAGTGGGAAGTGTACAACT |
| IDH2_418C>T | CAGACTCCAGAGCCCACACATTTGCACTCTAGACTCTACTGCCTTCCTCATGAAGAATTTTAGGACCCCCGTCTGGCTGTGTTGTTGCTTGGGGTTCAAAT<br>TCTGGTTGAAAGATGGCGGCTGCAGTGGGACCACTATTATCTCTGTCCTCACAGAGTTCAAGCTGAAGAAGATGTGGAAAAGTCCCAATGGAACTATCtGG<br>AACATCCTGGGGGGGACTGTCTTCCGGGAGCCCATCATCTGCAAAAACATCCCACGCCTAGTCCCTGGCTGGACCAAGCCCATCACCATTGGCAGGCACGC<br>CCATGGCGACCAGGTAGGCCAGGGTGGAGAGGGGATCCACTGACCTGGGCACCCCCCGACTGGAGCTCCTCGCCTAGCCATCCTCTTGTCTCTGCAGTACA<br>AGGCCACAGACTTTGTGGCAGACCGGGCCGGCACTTTCAAAATGGTCTTCACCCCAAAAGATGGCAGTGGTGTCAAGGAGTGGGAAGTGTACAACT |
| IDH2_419G>T | CAGACTCCAGAGCCCACACATTTGCACTCTAGACTCTACTGCCTTCCTCATGAAGAATTTTAGGACCCCCGTCTGGCTGTGTTGTTGCTTGGGGTTCAAAT<br>TCTGGTTGAAAGATGGCGGCTGCAGTGGGACCACTATTATCTCTGTCCTCACAGAGTTCAAGCTGAAGAAGATGTGGAAAAGTCCCAATGGAACTATCCtG<br>AACATCCTGGGGGGGACTGTCTTCCGGGAGCCCATCATCTGCAAAAACATCCCACGCCTAGTCCCTGGCTGGACCAAGCCCATCACCATTGGCAGGCACGC<br>CCATGGCGACCAGGTAGGCCAGGGTGGAGAGGGGATCCACTGACCTGGGCACCCCCCGACTGGAGCTCCTCGCCTAGCCATCCTCTTGTCTCTGCAGTACA<br>AGGCCACAGACTTTGTGGCAGACCGGGCCGGCACTTTCAAAATGGTCTTCACCCCAAAAGATGGCAGTGGTGTCAAGGAGTGGGAAGTGTACAACT |
| IDH2_419G>A | CAGACTCCAGAGCCCACACATTTGCACTCTAGACTCTACTGCCTTCCTCATGAAGAATTTTAGGACCCCCGTCTGGCTGTGTTGTTGCTTGGGGTTCAAAT<br>TCTGGTTGAAAGATGGCGGCTGCAGTGGGACCACTATTATCTCTGTCCTCACAGAGTTCAAGCTGAAGAAGATGTGGAAAAGTCCCAATGGAACTATCCaG<br>AACATCCTGGGGGGGACTGTCTTCCGGGAGCCCATCATCTGCAAAAACATCCCACGCCTAGTCCCTGGCTGGACCAAGCCCATCACCATTGGCAGGCACGC<br>CCATGGCGACCAGGTAGGCCAGGGTGGAGAGGGGATCCACTGACCTGGGCACCCCCCGACTGGAGCTCCTCGCCTAGCCATCCTCTTGTCTCTGCAGTACA<br>AGGCCACAGACTTTGTGGCAGACCGGGCCGGCACTTTCAAAATGGTCTTCACCCCAAAAGATGGCAGTGGTGTCAAGGAGTGGGAAGTGTACAACT |
| IDH2_515G>A | CAGACTCCAGAGCCCACACATTTGCACTCTAGACTCTACTGCCTTCCTCATGAAGAATTTTAGGACCCCCGTCTGGCTGTGTTGTTGCTTGGGGTTCAAAT<br>TCTGGTTGAAAGATGGCGGCTGCAGTGGGACCACTATTATCTCTGTCCTCACAGAGTTCAAGCTGAAGAAGATGTGGAAAAGTCCCAATGGAACTATCCGG<br>AACATCCTGGGGGGGACTGTCTTCCGGGAGCCCATCATCTGCAAAAACATCCCACGCCTAGTCCCTGGCTGGACCAAGCCCATCACCATTGGCAaGCACGC<br>CCATGGCGACCAGGTAGGCCAGGGTGGAGAGGGGATCCACTGACCTGGGCACCCCCCGACTGGAGCTCCTCGCCTAGCCATCCTCTTGTCTCTGCAGTACA<br>AGGCCACAGACTTTGTGGCAGACCGGGCCGGCACTTTCAAAATGGTCTTCACCCCAAAAGATGGCAGTGGTGTCAAGGAGTGGGAAGTGTACAACT |
| IDH2_515G>T | CAGACTCCAGAGCCCACACATTTGCACTCTAGACTCTACTGCCTTCCTCATGAAGAATTTTAGGACCCCCGTCTGGCTGTGTTGTTGCTTGGGGTTCAAAT<br>TCTGGTTGAAAGATGGCGGCTGCAGTGGGACCACTATTATCTCTGTCCTCACAGAGTTCAAGCTGAAGAAGATGTGGAAAAGTCCCAATGGAACTATCCGG<br>AACATCCTGGGGGGGACTGTCTTCCGGGAGCCCATCATCTGCAAAAACATCCCACGCCTAGTCCCTGGCTGGACCAAGCCCATCACCATTGGCAtGCACGC<br>CCATGGCGACCAGGTAGGCCAGGGTGGAGAGGGGATCCACTGACCTGGGCACCCCCCGACTGGAGCTCCTCGCCTAGCCATCCTCTTGTCTCTGCAGTACA<br>AGGCCACAGACTTTGTGGCAGACCGGGCCGGCACTTTCAAAATGGTCTTCACCCCAAAAGATGGCAGTGGTGTCAAGGAGTGGGAAGTGTACAACT |
| IDH2_516G>C | CAGACTCCAGAGCCCACACATTTGCACTCTAGACTCTACTGCCTTCCTCATGAAGAATTTTAGGACCCCCGTCTGGCTGTGTTGTTGCTTGGGGTTCAAAT<br>TCTGGTTGAAAGATGGCGGCTGCAGTGGGACCACTATTATCTCTGTCCTCACAGAGTTCAAGCTGAAGAAGATGTGGAAAAGTCCCAATGGAACTATCCGG<br>AACATCCTGGGGGGGACTGTCTTCCGGGAGCCCATCATCTGCAAAAACATCCCACGCCTAGTCCCTGGCTGGACCAAGCCCATCACCATTGGCAGcCACGC<br>CCATGGCGACCAGGTAGGCCAGGGTGGAGAGGGGATCCACTGACCTGGGCACCCCCCGACTGGAGCTCCTCGCCTAGCCATCCTCTTGTCTCTGCAGTACA<br>AGGCCACAGACTTTGTGGCAGACCGGGCCGGCACTTTCAAAATGGTCTTCACCCCAAAAGATGGCAGTGGTGTCAAGGAGTGGGAAGTGTACAACT |
| IDH2_516G>T | CAGACTCCAGAGCCCACACATTTGCACTCTAGACTCTACTGCCTTCCTCATGAAGAATTTTAGGACCCCCGTCTGGCTGTGTTGTTGCTTGGGGTTCAAAT<br>TCTGGTTGAAAGATGGCGGCTGCAGTGGGACCACTATTATCTCTGTCCTCACAGAGTTCAAGCTGAAGAAGATGTGGAAAAGTCCCAATGGAACTATCCGG<br>AACATCCTGGGGGGGACTGTCTTCCGGGAGCCCATCATCTGCAAAAACATCCCACGCCTAGTCCCTGGCTGGACCAAGCCCATCACCATTGGCAGtCACGC<br>CCATGGCGACCAGGTAGGCCAGGGTGGAGAGGGGATCCACTGACCTGGGCACCCCCCGACTGGAGCTCCTCGCCTAGCCATCCTCTTGTCTCTGCAGTACA<br>AGGCCACAGACTTTGTGGCAGACCGGGCCGGCACTTTCAAAATGGTCTTCACCCCAAAAGATGGCAGTGGTGTCAAGGAGTGGGAAGTGTACAACT |
| IDH2_514A>T | CAGACTCCAGAGCCCACACATTTGCACTCTAGACTCTACTGCCTTCCTCATGAAGAATTTTAGGACCCCCGTCTGGCTGTGTTGTTGCTTGGGGTTCAAAT<br>TCTGGTTGAAAGATGGCGGCTGCAGTGGGACCACTATTATCTCTGTCCTCACAGAGTTCAAGCTGAAGAAGATGTGGAAAAGTCCCAATGGAACTATCCGG<br>AACATCCTGGGGGGGACTGTCTTCCGGGAGCCCATCATCTGCAAAAACATCCCACGCCTAGTCCCTGGCTGGACCAAGCCCATCACCATTGGCtGGCACGC<br>CCATGGCGACCAGGTAGGCCAGGGTGGAGAGGGGATCCACTGACCTGGGCACCCCCCGACTGGAGCTCCTCGCCTAGCCATCCTCTTGTCTCTGCAGTACA<br>AGGCCACAGACTTTGTGGCAGACCGGGCCGGCACTTTCAAAATGGTCTTCACCCCAAAAGATGGCAGTGGTGTCAAGGAGTGGGAAGTGTACAACT |
| IDH2_514A>G | CAGACTCCAGAGCCCACACATTTGCACTCTAGACTCTACTGCCTTCCTCATGAAGAATTTTAGGACCCCCGTCTGGCTGTGTTGTTGCTTGGGGTTCAAAT<br>TCTGGTTGAAAGATGGCGGCTGCAGTGGGACCACTATTATCTCTGTCCTCACAGAGTTCAAGCTGAAGAAGATGTGGAAAAGTCCCAATGGAACTATCCGG<br>AACATCCTGGGGGGGACTGTCTTCCGGGAGCCCATCATCTGCAAAAACATCCCACGCCTAGTCCCTGGCTGGACCAAGCCCATCACCATTGGCgGGCACGC<br>CCATGGCGACCAGGTAGGCCAGGGTGGAGAGGGGATCCACTGACCTGGGCACCCCCCGACTGGAGCTCCTCGCCTAGCCATCCTCTTGTCTCTGCAGTACA<br>AGGCCACAGACTTTGTGGCAGACCGGGCCGGCACTTTCAAAATGGTCTTCACCCCAAAAGATGGCAGTGGTGTCAAGGAGTGGGAAGTGTACAACT |

**Table S28. Sequence of gBlocks used in the As-BDA *IDH2* assay.**

**REFERENCE**

1. Tate, J. G. *et al.* COSMIC: the Catalogue Of Somatic Mutations In Cancer. *Nucleic Acids Res.* **47**, D941–D947 (2019).
2. Song, P. *et al.* Selective multiplexed enrichment for the detection and quantitation of low-fraction DNA variants via low-depth sequencing. *Nat. Biomed. Eng.* (2021) doi:10.1038/s41551-021-00713-0.
3. Cheng, L. Y. *et al.* High sensitivity sanger sequencing detection of BRAF mutations in metastatic melanoma FFPE tissue specimens. *Sci. Rep.* **11**, 9043 (2021).
